# Multimodal Mapping of Human Lymphopoiesis Reveals B and T/NK/ILC Lineages are Subjected to Cell-Intrinsic *versus* Flt3L-Dependent Regulation

**DOI:** 10.1101/2022.12.12.520022

**Authors:** Kutaiba Alhaj Hussen, Emna Chabaane, Elisabeth Nelson, Shalva Lekiashvili, Samuel Diop, Seydou Keita, Bertrand Evrard, Aurélie Lardenois, Marc Delord, Els Verhoeyen, Kerstin Cornils, Zeinab Kasraian, Elizabeth A. Macintyre, Ana Cumano, David Garrick, Michele Goodhardt, Guillaume P. Andrieu, Vahid Asnafi, Frederic Chalmel, Bruno Canque

## Abstract

The developmental cartography of human lymphopoiesis remains incompletely understood. Here, we establish a multimodal map that extends the current view of lymphoid development. Our results demonstrate that lymphoid specification follows independent direct or stepwise differentiation pathways converging toward the emergence of CD117^lo^ multi-lymphoid progenitors (MLPs) that undergo a proliferation arrest before entering the CD127^-^ (T/NK/ILC) or CD127^+^ (B) lymphoid pathways. While the emergence of CD127^-^ early lymphoid progenitors is driven by Flt3 signaling, differentiation of their CD127^+^ counterparts is regulated cell-intrinsically and depends exclusively on the divisional history of their precursors. Single-cell mapping of lymphoid differentiation trajectories reveals that a dissociation between proliferation and differentiation phases allows amplification of the precursor pools prior to the onset of antigen receptor rearrangement. Besides demonstrating that B and T/NK/ILC lineages are subjected to differential cell-autonomous *versus* Flt3-inducible regulation, our results go a long way to reconciling human and mouse models of lymphoid architecture.

## INTRODUCTION

Hematopoiesis is defined as the physiological process by which multipotent, lineage-specified stem/progenitor and precursor cells self-renew or differentiate along divergent pathways to ensure the lifelong production of platelets, red blood cells and leukocytes. Over the last decade, single-cell genomics and fate mapping studies have revealed a previously unsuspected functional heterogeneity within the hematopoietic stem/progenitor cell (HSPC) compartment. Together these studies suggest that rather than as a stepwise hierarchy comprising distinct populations, hematopoiesis is organized as a developmental continuum where hematopoietic stem cells (HSCs) feed a continuous flux of differentiation along the different lineage branches ^1^. As well as passing through intermediate stages of multipotent progenitors, it is now clear that HSCs can directly commit toward the erythroid or megakaryocytic lineages ^2–4^ or enter the granulocytic or monocytic lineages in response to myeloid growth factors ^5, 6^. Consistent with this, clonal scale in vivo tracking of HSC differentiation under steady-state or regenerative conditions in mouse ^7, 8^ or non-human primate ^9^ models, as well as transplanted patients ^10, 11^, provided evidence that lineage priming can be initiated at the level of HSCs. Despite this, there is still debate regarding when and how progenitors execute fate decisions and if lineage choices are primarily conditioned by intrinsic cell properties, stochastic behavior or dictated by extracellular signals ^12^. Further, increasing evidence indicates that lymphoid and myeloid lineages are subjected to distinct developmental constraints. This is supported by transplantation studies showing that the early emergence of myeloid cells contrasts with the delayed lymphoid reconstitution ^13, 14^, as well as by previous reports in the mouse that lymphoid specification initiates downstream of HSCs, at the level of lymphoid-primed multipotent progenitors (LMPPs) ^15^. LMPPs critically depend on Flt3 signaling for survival and expansion ^16^ and stepwise progression to downstream RAG1- or DNTT-expressing early or lymphoid-primed progenitors (ELPs/LPPs) ^17–19^, and then to IL7R/CD127^+^ common lymphoid progenitors (CLPs) ^20^.

The developmental architecture of human lymphopoiesis remains less well characterized than in the mouse. It is nonetheless established that upregulation of CD45RA by CD34^hi^ hematopoietic stem/progenitor cells (HSPCs) marks the loss of erythro-megakaryocytic potential ^21, 22^ and that lymphoid fate co-segregates with granulomonocytic and dendritic potentials within the CD45RA^+^ compartment ^23–25^. Earlier studies also identified candidate counterparts of mouse CLPs, usually referred to as the CD7 ^25, 26^ or CD10 ^23, 27, 28^ multi-lymphoid progenitors (MLPs) whose functions and developmental statuses remain controversial ^29^. Similarly, no human counterparts of murine LMPPs have been formally identified to date ^28^. To overcome constraints inherent to studies of human hematopoiesis, we have previously developed an in-vivo modeling approach of mid-fetal hematopoiesis in humanized mice ^30^. This led to the demonstration that human lymphopoiesis displays a bipartite organization stemming from founder populations of CD127^-^ and CD127^+^ ELPs and that, whereas the CD127^-^ ELPs are mainly T/NK/ILC precursors, their CD127^+^ counterparts are intrinsically biased toward the B lineage ^31^. However, at present the precise origin of these ELPs, and the regulation of their developmental relationships are still unknown.

In the current study, combining time-course and endpoint molecular and functional analyses, we demonstrate that CD127^-^ and CD127^+^ ELPs originate from a previously unidentified subset of CD117^lo^ MLPs and that they are subjected to a differential cell-intrinsic versus Flt3L-dependent regulation.

## RESULTS

### CD127^-^ and CD127^+^ ELP production patterns depend on the divisional history of their precursors

We analyzed hematopoietic reconstitution in mice xenografted with neonatal CD45RA^-^Lin^-^ HSCs or CD45RA^hi^Lin^-^ LMDPs (Lympho-Myelo-Dendritic Progenitors) ^31^ transduced with *egfp*-reporter lentiviruses (Fig. 1A and Extended Data Fig. 1A, B). Dynamic follow-up of HSC-engrafted mice confirmed that reconstitution of myeloid monocytic or dendritic cells precedes that of B lymphocytes and disclosed a sequential emergence of CD127^-^ and CD127^+^ ELPs (Fig. 1B, C). This also showed that expansion of the B cell compartment closely parallels with the increase in the production of CD127^+^ ELPs. Overall similar, albeit accelerated, biphasic reconstitution patterns were observed in LMDP-engrafted mice that supported only transient hematopoietic reconstitution (Fig. 1D, E). To investigate whether sequential emergence of CD127^-^ then CD127^+^ ELPs reflects a temporal shift in the lymphoid potential, we developed an in vitro diversification assay in which myeloid granulocyte (Gr; CD15^+^), monocyte (Mo; CD115^+^) and dendritic (DC; CD123^+^) precursors, as well as lymphoid CD127^-^ or CD127^+^ ELPs and CD19^+^ B lymphocytes are quantified as the readout of differentiation potentials (Extended Data Fig. 2A). Therefore, LMDPs were sorted at weekly endpoints from HSC-engrafted mice and seeded for 7 or 14 days onto OP9 stroma under low serum conditions supplemented with SCF (hereafter referred to as the standard condition) (Fig. 2A). FACS analyses at day-7 revealed that, in contrast to the stable production of monocyte and dendritic cells, lymphoid outputs follow an ascendant trend driven by the gradual increase in CD127^+^ ELP production levels (Fig. 2B). Consistent with these observations, a concordant increase in CD19^+^ BLs was noted in cultures at day-14 (Fig. 2C). Subsequent finding that the lymphoid potential of LMDPs sorted at weekly endpoints from LMDP-engrafted mice evolves in the same way reinforced the view that the time-dependent increase in CD127^+^ ELP production levels reflect cell-intrinsic variations in the lymphoid potential of their precursors. (Extended Data Fig. 2B-F).

**Figure 1.**
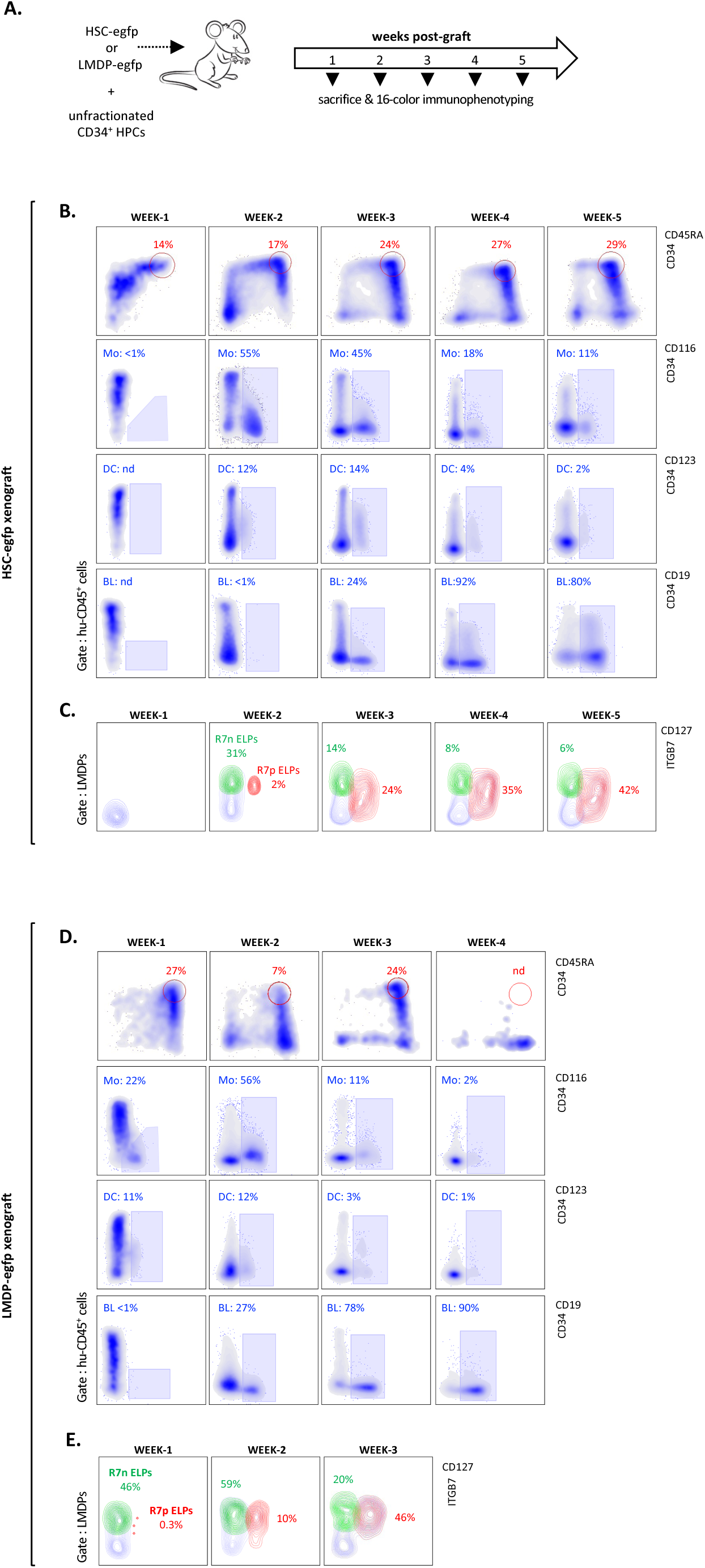
Dynamics of hematopoietic reconstitution in xenografted mice. (A). Experimental design: neonatal UCB HSCs or LMDPs transduced with egfp-reporter lentiviruses (1.5 x 10^5^ cells/mouse) were mixed with non-transduced bulk CD34^+^ HPCs (1.5 x 10^5^ cells/mouse) before intravenous injection to irradiated NSG mice. To follow multi-lineage reconstitution, hu-CD45^+^egfp^+^ cells harvested from the BM of mice sacrificed at weekly endpoints were analyzed by multiparameter flow cytometry. (B, C). Dynamic follow up of mice reconstituted with lentivirally-transduced HSCs. (B) Density plots show the kinetic of BM CD34^hi^CD45RA^hi^ HSPCs (upper panels; red circles), monocytes (Mo; CD116^+^CD123^-^) (medium panels; blue rectangles), dendritic cells (DC; CD116^+^CD123^+^) and B lymphocytes (lower panels; blue rectangles) in mice reconstituted with lentivirally-transduced HSCs (gates: hu-CD45^+^egfp^+^ cells). (C) Contour plots show the sequential emergence of CD127^-^ (green) and CD127^+^ (red) ELPs; gates: huCD45^+^CD34^+^CD45RA^hi^Lin^-^egfp^+^ HSPCs. Percentages are indicated; results from individual mice are representative of 3 independent experiments. (D-E). Dynamic follow up of mice reconstituted with lentivirally-transduced LMPS. Bidimensional density and contour plots show the dynamics of same populations as above. Results from individual mice are representative of 2 independent experiments.

**Figure 2.**
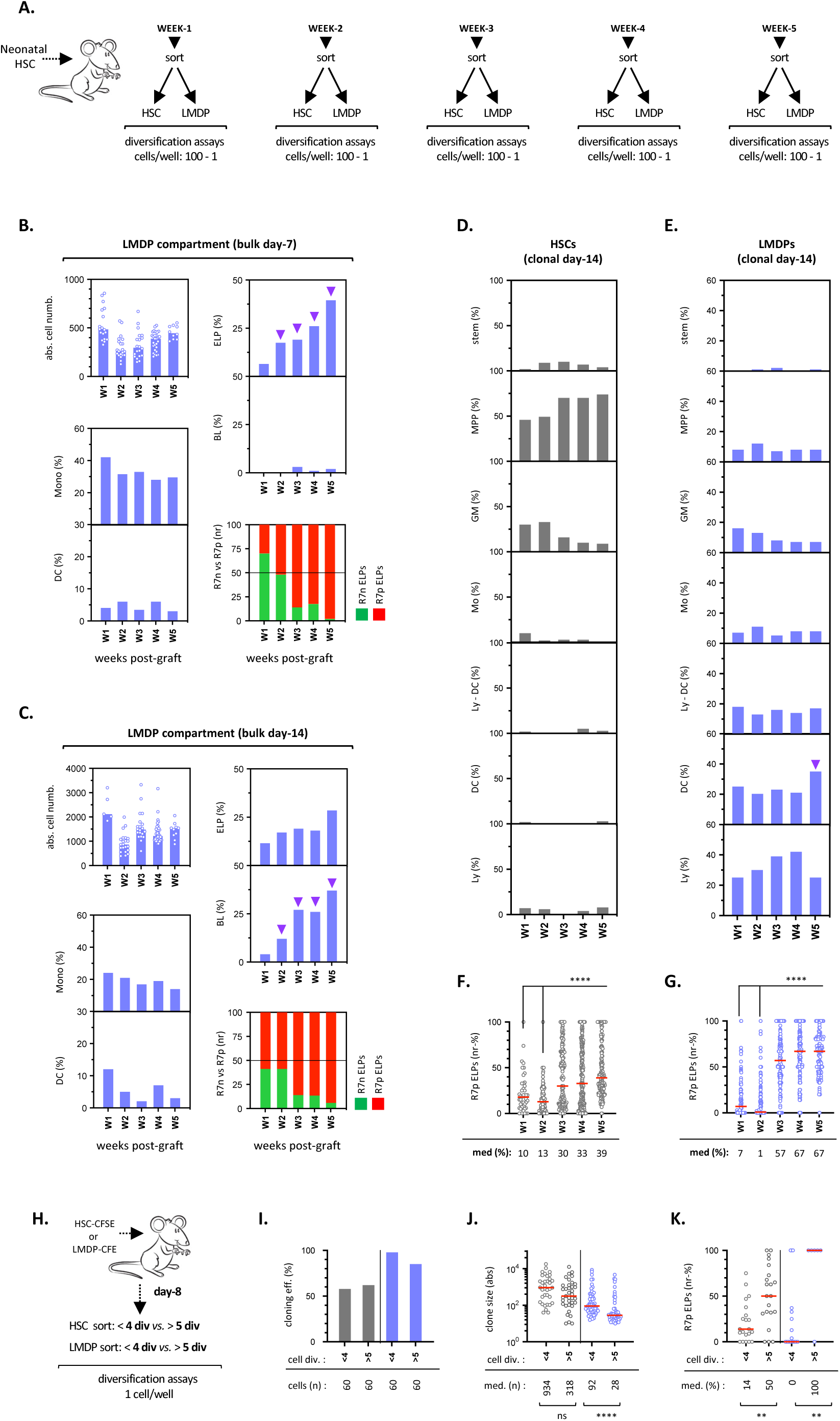
In vitro assessment of lymphoid potential in multi-lineage diversification assays. (A). Experimental design: mice xenografted neonatal HSCs were sacrificed at the indicated endpoints before assessment of HSC or LMDP lympho-myeloid potential in bulk or single-cell in vitro diversification assays. (B, C). Assessment of lineage output in bulk diversification assays. LMDPs were cultured by 100 cell-pools for (B) 7 or (C) 14 days under standard *scf* condition before quantification of absolute cell yields and lineage outputs. Upper left panel: absolute cell yields (bars indicate median cell numbers; circles correspond to individual wells); lower left panels: median percentages of CD115^+^ monocytes (top) and CD123^+^ DC (bottom); upper right panels: median percentages of CD127^-/+^ ELPs (top) and BLs (bottom); results are normalized relative to total hu-CD45^+^ cells; lower right stacked bar plot: normalized ratios of CD127^-^ (green) *vs*. CD127^+^ (red) ELPs. Results expressed as median percentages of 5 to ≥ 20 replicates are from one representative experiment out of 3; purple arrows show the time-dependent increase in ELP or B-cell outputs. (D-G). Assessment of lineage output in clonal diversification assays. HSC or LMDPs were cultured for 14 days under *scf:gm-csf:tpo* condition before quantification of absolute cell yields and lineage outputs; positivity threshold for clone detection was set arbitrarily at ≥ 20 cells/clone. (D, E) Bar plots show the dynamics of Stem (Stem: CD34^+^ cells ≥50%; others <10%), multipotent (MPP: Gr ≥ 10%; Mono ≥ 10%; DC ≥ 10%; Ly ≥ 10%), granulomonocytic (GM: Gr + Mono ≥50%; others <10%), monocytic (M: Mono ≥50%; others <10%), lympho-dendritic (Ly-DC: Ly + DC ≥50%; others <10%), dendritic (DC: ≥50%; others <10%) or lymphoid (Ly: Ly: ≥50%; others <10%) clones across the (D) HSC (gray bars) or (E) LMDP (blue bars) compartments; purple arrow shows the increase in DC progenitors observed at wek-5 after grafting. (F, G) Circle plots show the kinetics of CD127^+^ ELPs within lymphoid-containing (F) HSC- or (G) LMDP-derived clones; results are normalized relative to total ELP content and expressed on a per clone basis. Red bars indicate medians; corresponding values are indicated (lower row); positivity threshold for ELP detection is set arbitrarily at ≥ 10 ELPs/clone). Assessment of statistical significance was performed with the Mann-Whitney test (**** p<0.0001; *** p<0.001; ** p<0.01). Results are pooled from 2 experiments. (H-K). Effect of conservative cell divisions on the lymphoid potential of HSC or LMDPs. (H) Experimental design: neonatal HSCs or LMDP were labeled with proliferation-dependent dye carboxyfluorescein diacetate succinimidyl ester (CFSE) prior to injection to irradiated NSG mice (2.5 x 10^5^ cells/mouse). At day 8 after grafting, self-renewed HSCs or LMDPs subdivided according to the number of conservative cell divisions (<4 *versus* >5 divisions) were seeded by FACS in single-cell diversification assays. (I) Cloning efficiencies of divided (>5) *versus* non-divided (<4) HSCs (gray bars) or LMDPs (blue bars); (J) Absolute cell numbers per clone derived from divided *versus* non-divided HSCs (gray circles) or LMDPs (blue circles); (K) Relative percentages of CD127^+^ ELPs detected among lymphoid-containing clones derived from divided *versus* non-divided HSCs (gray circles) or LMDPs (blue circles). Red bars indicate medians; corresponding values are indicated (lower row). Assessment of statistical significance was performed as above.

To further document this point, we optimized the diversification assay to read out the lympho-myeloid potential at a clonal resolution. We found that adding GM-CSF and TPO to SCF results in high cloning efficiency and allows multilineage diversification. Flow cytometry analysis of clones derived from single HSCs or LMDPs sorted as above at weekly endpoints from HSC-engrafted mice showed that cell cloning efficiencies and expansion rates remain stable throughout the 5-week follow-up period (Extended Data Fig. 3A-D). Stratification of clones according to lineage output confirmed that the HSC compartment (Fig. 2D) comprises a majority of multipotent progenitors (MPPs) and revealed that granulomonocytic progenitors (GM) reach maximum levels during the first 2-weeks after grafting. Analysis of downstream LMDPs (Fig. 2E) disclosed the expected enrichment in bi- or uni-lineage progenitors and found that, except for a late increase in dendritic progenitors, their clonal architecture remains stable over time. Applying the Uniform Manifold Approximation and Projection (UMAP) algorithm on a compendium of 2,806 clones showed that the distribution of HSC or LMDP ancestor cells is primarily driven by expansion rates with highly proliferative multipotent ancestors projecting at the root of the map followed by sequential branching of granulomonocytic, dendritic and then lymphoid precursors (Extended Data Fig. 3E-G). Hierarchical clustering according to differentiation potentials confirmed the late increase in dendritic progenitors amongst LMDPs (Extended Data Fig. 3H). Again, focusing on ELP output (Fig. 2F, G) revealed that, irrespective of their HSC or LMDP origin and, most importantly, independent of overall lymphoid potential, lymphoid-containing clones displayed increasing CD127^+^ ELP output beyond the 2^nd^ week after grafting, consistent with the above results.

To examine whether time-dependent variation in CD127^-^ and CD127^+^ ELP production pattern depends on the division history of their precursors, HSCs or LMDPs isolated at day 8 from mice reconstituted with CFSE-labelled HSCs or LMDPs were fractionated according to the number of conservative divisions they had undergone (<4 versus > 5 divisions) and seeded as above in clonal assays (Fig. 2H and Extended Data Fig. 4A, B). Quantification of ELP output confirmed that HSC or LMDPs which had undergone a higher number of previous cell divisions show a marked bias towards the production of CD127^+^ ELPs (Fig. 2I-K).

Collectively, these results indicate that the rise in CD127^+^ ELP production over time reflects qualitative changes in the lymphoid potential of their precursors.

### CD127^-^ and CD127^+^ ELPs differentiate from bipotent MLPs

Our observation that LMDPs comprise lymphoid-restricted progenitors led us to prospectively isolate candidate MLPs. Further immunophenotypic stratification revealed that decreasing CD117 expression levels coincides with downmodulation of myeloid CD33 (Extended Data Fig. 4C). To test whether lymphoid specification can be delineated on these bases, LMDPs were subdivided into three CD117^hi-int-lo^ fractions before seeding in bulk diversification assays with graded doses of SCF (Fig. 3A-C). Analysis of lineage output confirmed that downmodulation of CD117 correlates with the rise in lymphoid potential as well as concomitant decline in the cell expansion rates. This also found that, independently of overall lymphoid production and CD117 expression levels, low doses of SCF skew lymphoid differentiation toward the CD127^+^ ELPs and the B lineage (Fig. 3D). Further assessment of lympho-myeloid potentials in cultures supplemented with lymphoid or myeloid growth factors (Extended Data Fig. 4D-F) showed that CD117^hi^ LMDPs expand vigorously in response to G-CSF, indicating enrichment in granulomonocytic progenitors, whereas CD117^int^ LMDPs display mixed lympho-dendritic. Again, irrespective of culture conditions, the CD117^lo^ LMDP fraction showed lymphoid-restricted output. With respect to ELP production patterns, this also revealed that adding TPO and/or GM-CSF resulted in balanced ELP outputs while supplementation in Flt3L induces a complete shift toward the CD127^-^ subset (Extended Data Fig. 4G). In addition, irrespective of culture conditions, CD117^lo^ LMDPs failed to expand in vitro which is indicative of an intrinsically low proliferative capacity.

**Figure 3.**
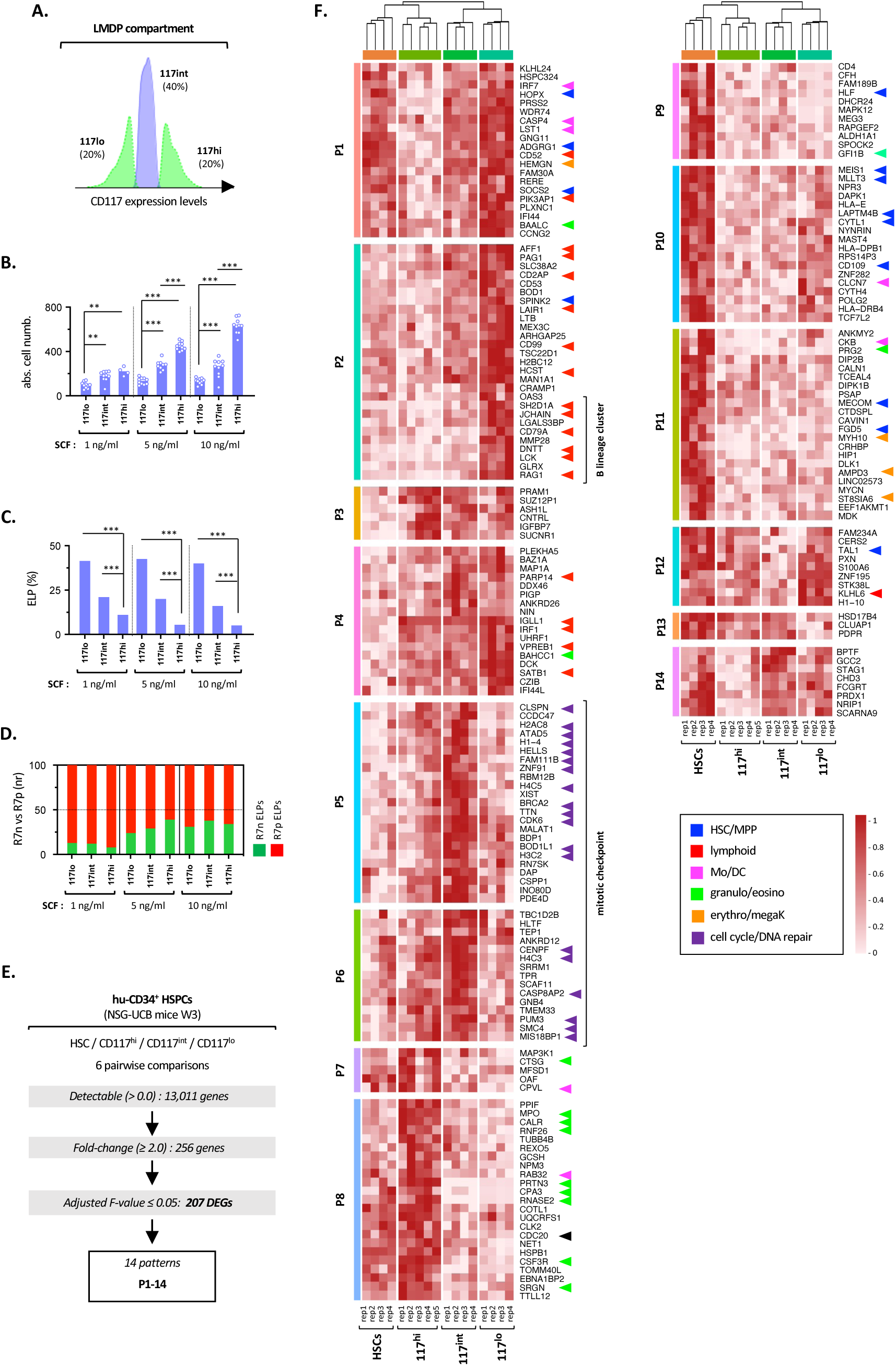
Molecular and functional characterization of CD117^lo^ MLPs. (A). Immunophenotypic fractionation of LMDPs based on CD117 expression levels. The full gating procedure in shown in Extended Data Fig.1A. (B-D). In vitro assessment of the lymphoid potential of CD117^high, int, low^ LMDPs. The indicated subsets were sorted at week-3 from the BM of HSC-xenografted mice and seeded by 100 cell/pools for 7 days in diversification assays supplemented with increasing doses of SCF. (B) Absolute cell yields; bars indicate median cell number; circles correspond to individual wells; (C) Percentages of CD127^-/+^ ELPs; results are normalized relative to total hu-CD45^+^ cells; bars indicate medians; (D) Stacked bar plots show normalized ratios of CD127^-^ (green) *vs.* CD127^+^ (red) ELPs. Results are expressed as median percentages of 15 replicates from one representative experiment. Assessment of statistical significance was performed with the Mann-Whitney test (**** p<0.0001; *** p<0.001; ** p<0.01). (E-G). Transcriptional profiling of HSC and CD117^high, int, low^ LMDP fractions. (E) Flowchart summarizing the filtration and clustering strategies used for extraction of population-specific gene signatures across the corresponding cellular subsets; detection thresholds, fold-changes and adjusted F-value are indicated. (F) Heatmap showing expression of 207 selected DEGs across 14 clusters; gene expression values are log-transformed, normalized, and scaled; colored arrows show lineage- or subset-specific genes.

To substantiate these findings, CD117^hi-int-lo^ LMDP fractions, as well as HSCs, were subjected to transcriptional profiling with an ultra-low-input mini-RNA-seq method. Pairwise comparisons followed by unsupervised gene and population clustering identified 207 differentially expressed genes (DEG) partitioned into 14 gene expression patterns (P1-14) (Fig. 3E and Table S1A, B). Functional analysis and pattern distribution (Fig. 3F) confirmed that HSCs selectively express genes regulating quiescence (P1, P12: *SOCS2, TAL1, HOPX*) ^32, 33^ or self-renewal (P9-10: *HLF*, *MLLT3*, *MEIS1*) ^34^, as well as early regulators of erythro-megakaryocytic differentiation (P11: *AMPD3*, *MYH10*). Consistent with enrichment in granulomonocytic potential, CD117^hi^ LMDPs overexpressed *CSF3R*, *MPO*, *PRTN3* and *CTSG* (P7-8). As expected, decreasing CD117 expression coincided with extinction of granulomonocytic lineage genes and concomitant upregulation of early lymphoid regulators (P2-4: *AFF1*, *SATB1*, *IRF1*) and a series of lymphoid marker genes (P2-4: *LAIR1*, *CD2AP*, *IGLL1*, *HSCT*, *VPREB1*) whose expression levels peaked in the CD117^lo^ fraction. Consistent with their intrinsic B-lineage bias, lymphoid-committed CD117^lo^ LMDPs displayed further overexpression of *LCK*, *JCHAIN*, *CD79A*, *DNTT* and *RAG1* transcripts (P2, bottom). Genes involved in cell-cycle-related pathways (P5-6: DNA replication/repair, G2M mitotic checkpoint) were also enriched in CD117^int^ LMDPs, consistent with their active proliferation. Expression of these genes dramatically decreased in downstream CD117^lo^ compartment, consistent with their low expansion rates. Importantly, CD117^lo^ LMDPs also retained (or upregulated) a repertoire of stem cell-associated genes (P1-10-12: *MEIS1*, *MLLT3, TAL1*, *HOPX, SOCS2*).

Altogether, these results demonstrate that CD117^lo^ LMDPs correspond to prototypic MLPs. They also provide evidence that lymphoid commitment is associated with a proliferation arrest.

### Flt3 signaling regulates NK/ILC/T-*versus* B-lineage choice at the level of CD117^lo^ MLPs

To further investigate the signaling pathways regulating the choice between CD127^-^ and CD127^+^ ELPs, CD117^lo^ MLPs were subsequently cultured onto OP9 or OP9-Dl4 stroma with SCF and a series of 15 soluble ligands corresponding to growth/differentiation factors and pro- or anti-inflammatory cytokines (Table S2A). Whereas adding TNF-*a*, TGF-*b*1 or retinoic acid (RA) inhibited ELP differentiation, supplementation in Flt3L or IL-7 led to an almost complete disappearance of the CD127^+^ subset (Fig. 4A). Notch1 signaling also favored CD127^-^ ELP differentiation from which a majority upregulated surface CD7 indicative of T-lineage commitment (Fig. 4B and data not shown). Analysis by multiplex PCR of in vitro-derived ELPs (Extended Data Fig. 5 and Table S2B) disclosed the expected fingerprints ^31^ within CD127^-^ (*MEF2C*, *RUNX2*, *IKZF1*, *ID2*, *NFIL3*) and CD127^+^ (*IL7R*, *RAG2*, *EBF1*, *FBXW7*) ELPs generated under SCF, TSLP or Flt3L conditions, and confirmed that a 7-day culture on OP9-Dl4 stroma drives the emergence of prototypic T-cell precursors (*DTX1*, *NOTCH3*, *LEF1*, *TOX* and *CCR9*). Notably, CD127^-^ ELPs generated under the IL-7 condition downmodulated most transcripts, precluding further lineage affiliation. To screen for candidate inducers of CD127^+^ ELP differentiation, we tested the effect of 18 ligands of receptors whose transcripts are differentially expressed between LMDPs and the CD127^-/+^ ELPs ^31^ (Table S2B) but here again with essentially negative results. This reinforces the idea that the differentiation of CD127^+^ ELPs depends on cell-autonomous regulatory mechanisms.

**Figure 4.**
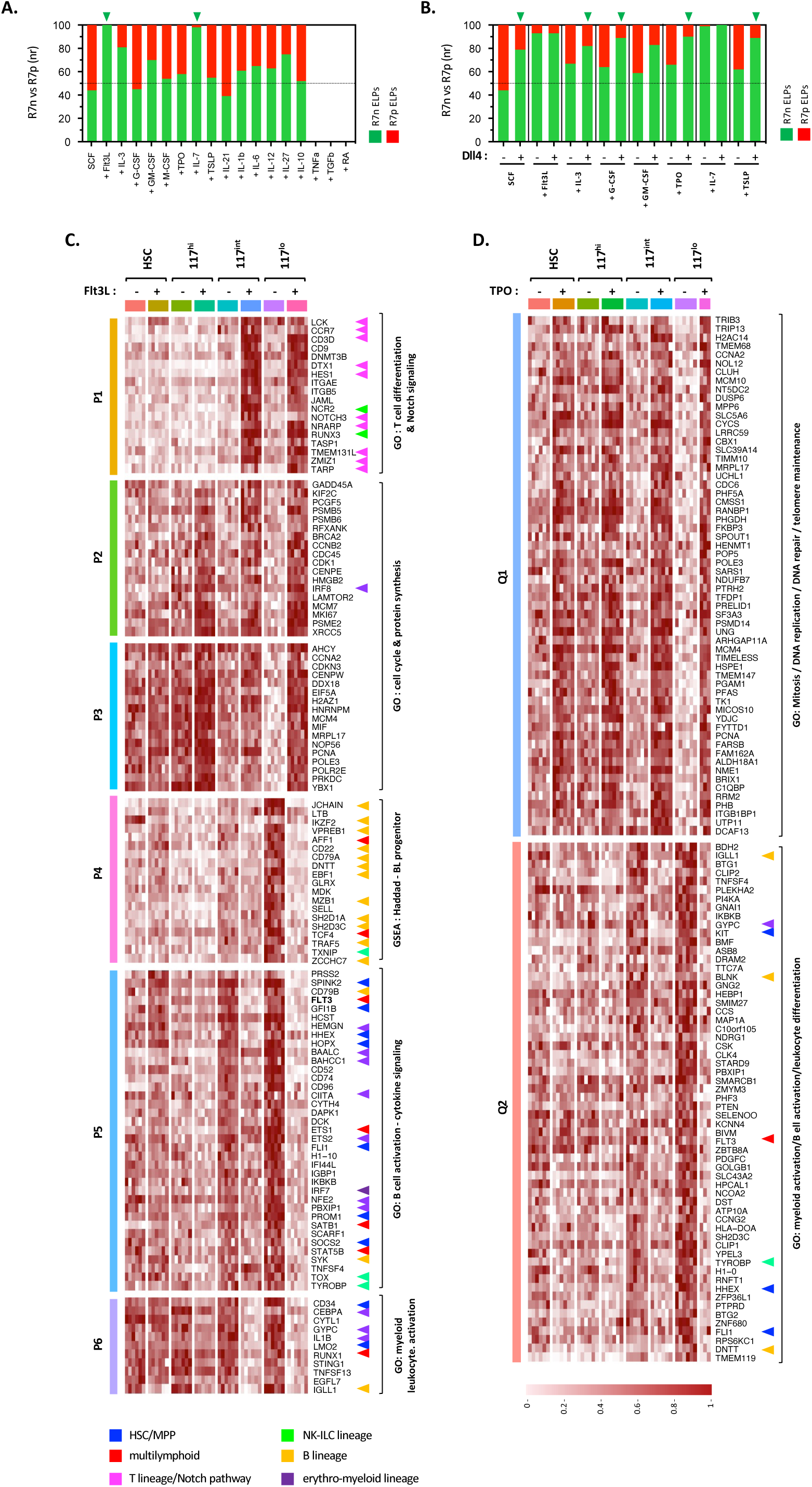
Effect of soluble factors on T/NK/ILC versus B lineage choice. (A, B). MLPs isolated as above of the BM of HSC-xenografted mice seeded by 100 cell-pools onto (A) OP9 or (B) OP9-DL4 stroma were cultured for 7 days under standard *scf* condition supplemented with the indicated ligands (all used at 1 ng/ml, except for RA: 10^-5^ M) before quantification of CD127^-^ and CD127^+^ ELP output. Stacked bar plots show normalized ratios of CD127^-^ (R7n; green) *vs.* CD127^+^ (R7p; red) ELPs; results are expressed as median percentages of 5 to ≥ 20 replicates pooled from independent experiments. (C, D) Transcriptional response of CD117^lo^ MLPs to short-term stimulation with Flt3L or TPO. The indicated HSC or CD117^hi, int^ LMDP or CD117^lo^ MLP subsets were seeded by 500 cell pools in 96-well plates and cultured for 6 hours under SCF condition with or without Flt3L or TPO (10 ng/ml each) before being processed for mini-RNAseq. Heatmaps show expression of compendia of 120 (A) Flt3L- or (B) TPO-responsive genes within the indicated populations; gene expression values are log-transformed, normalized, and scaled; genes linked to Notch pathway, multi-lymphoid progenitors (blue) or T/NK- (green) or B- (red) or NK/ILC- (yellow) lineages are indicated.

To get insight into the gene regulatory networks controlling the dichotomic choice between CD127^-^ and CD127^+^ ELPs, CD117^lo^ MLPs, as well as CD117^hi-int^ LMDPs and CD45RA^-^Lin^-^ HSCs were seeded for 6 hours under the standard SCF condition with or without Flt3L or TPO before analysis by mini-RNA-seq. Gene expression analyses based on global pairwise comparisons allowed selection of a compendium of 972 DEGs among which 638 Flt3L- and 544 TPO-responsive genes were identified (Extended Data Fig. 6A). Side-by-side comparison between treated or untreated cellular subsets revealed that CD117^lo^ MLPs are the most responsive to Flt3L with 454 DEGs, as compared to 193 for CD117^int^ LMDPs and <80 for the other HSC or CD117^hi^ LMDP subsets (Extended Data Fig. 6B and Tables S3A, B). Functional analysis of DEGs confirmed that in CD117^lo^ MLPs and to a lesser extent CD117^int^ LMDPs (Fig. 4C), Flt3L induces a major developmental transition which is characterized by upregulation of genes linked to NK/ILC/T differentiation (*LCK*, *NCR2*, *CD3*, *TARP*, *TMEM131L*, *CCR7*) or Notch signaling (*DTX1*, *HES1*, *Notch3*, *NRARP*) (P1), as well as of a broad repertoire of genes involved in cell cycle, DNA repair and protein synthesis pathways (P2-3). Concomitantly, genes controlling early lymphoid development (*AFF1*, *SATB1*, *RUNX1*) or B-lineage specification (*IKZF2*, *ETS1*, *EBF1*, *VPREB1*, *DNTT*, *JCHAIN*) (P4), as well as genes associated with stem/progenitor cell maintenance (*SOCS2*, *HOPX*, *LMO2*) or erythro-myeloid differentiation (*NFE2*, *GYPC*, *ETS2*, *GFI1B*, *CEBPA, IRF7*) (P5-6) were subjected to an active repression. Notably, Flt3L-treated cells also downmodulated *FLT3* transcripts, suggesting a potential negative auto-regulatory loop. In contrast to the effect of Flt3L but consistent with its established role in stem cell maintenance ^35^, short-term stimulation with TPO resulted in upregulation of a wide repertoire of genes involved in DNA repair, telomere maintenance or cell cycle regulation (Q1) and corresponding downmodulation of *KIT* and *FLT3*, and diverse lineage-specific marker genes (Q2: *IGLL1*, *BLNK*, *DNTT*, *GYPC*, *TYROBP*) (Fig. 4D and Tables S4A, B). Conversely to Flt3L, only minor differences in population-specific transcriptional responses were noted between TPO-treated subsets (Extended Data Fig. 6C).

These results demonstrate that Flt3L represses the intrinsic B-lineage bias of CD117^lo^ MLPs to regulate the dichotomic choice between CD127^-^ and CD127^+^ lymphoid pathways.

### Increasing expression of B-lymphoid genes drives the dynamics of regenerative lymphopoiesis

To further investigate the mechanisms regulating CD127^+^ ELP differentiation, we next searched for molecular correlates of the temporal variations in lymphoid production patterns described above. Therefore, CD117^lo^ MLPs, as well as CD117^hi-int^ LMDPs and HSCs, were sorted at weekly interval from the BM of HSC-xenografted mice and analyzed by mini-RNA-seq (Fig. 5A). Pairwise comparisons across all time points and cell populations (76 pairwise comparisons) revealed a compendium of 2207 genes subjected to time-dependent variation and showed that they distribute in 5 predominant expression patterns (P1-5) (Extended Data Fig. 6D, E). Functional enrichment showed that the expression level of genes controlling cell proliferation, protein synthesis or ATP metabolic processes peaks at week-1 after grafting (P1) to gradually decrease thereafter whereas genes involved in interferon signaling pathways (P2-4) follow an opposite upwards trend (Extended Data Fig. 7A-D and Tables S5-8). Consistent with the late expansion of DC progenitors detected above, cell populations isolated from mice at week-5 after grafting overexpressed transcripts coding for regulators of DC differentiation (*IRF7*, *IRF8*, *SPIB*, *TCF4*) (P3-5). As expected, these analyses also revealed that over time HSCs upregulate a set of B-lymphoid genes (*BCL11A*, *IKZF3*, *VPREB1*, *JCHAIN*, *IGLL5*, *CD79B*) correlating with the B-lineage polarization of their lymphoid output (Extended Data Fig. 7A). Consistent with this observation, the repertoire of B-lymphoid genes subjected to time-dependent deregulation broadened as cells progressed along the lymphoid specification axis, reaching maximum diversity and expression levels in CD117^lo^ MLPs isolated from mice at week-5 after grafting (Extended Data Fig. 7D and Table S8). At that time, CD117^lo^ MLPs expressed at the highest levels *SATB1*, *MEF2A*, *EBF1*, *LEF1* and *SOX4* TFs, as well as genes linked to IL-7 signaling (IL*-7R*, *JAK3*) and antigen receptor rearrangement (*ADA*, *DNTT*, *RAG1*), which here again was indicative of increasing specification toward the B lineage. Dynamic follow-up of 20 lymphoid genes across all time points and cell compartments (Fig. 5B) confirmed that over time the expression levels of genes linked to B-cell differentiation (*EBF1*, *BCL11A*), IL7R (*IL7R*, *JAK3*) or pre-BCR signaling (*CD79A*, *VPREB1*, *BLNK*) and TCR/Ig rearrangement (*RAG1*, *DNTT*) follow parallel increasing trends. In contrast to *CD3D* whose expression levels declined over time, *FLT3* transcript levels followed similar upward trend which suggests that despite an increasing B-lineage bias CD117^lo^ MLPs remain fully responsive to Flt3L.

**Figure 5.**
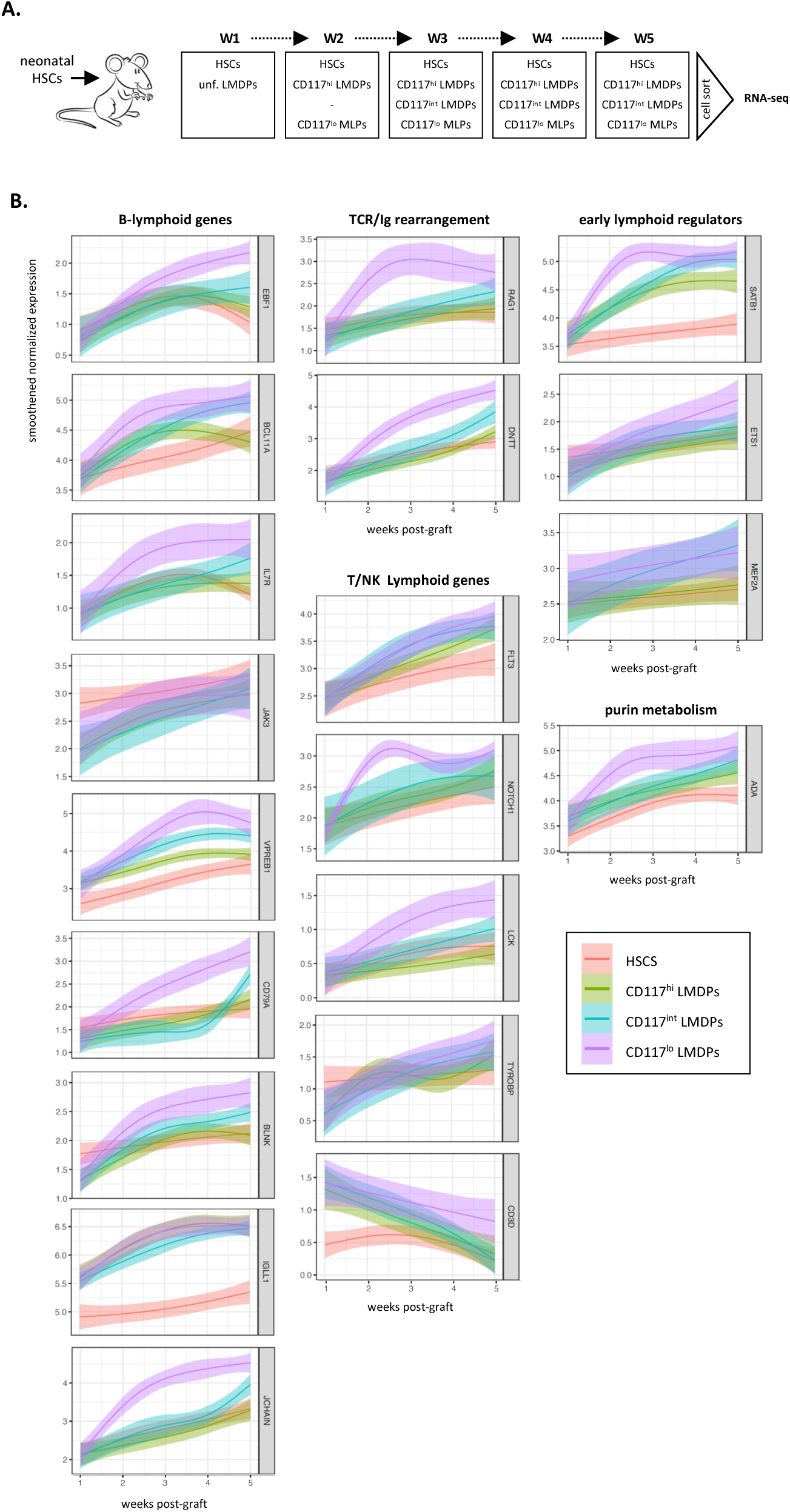
Temporal variations in expression of 20 lymphoid genes across 4 HSPC subsets. (A). Experimental design: HSC-xenografted mice were sacrificed at the indicated time points before the indicated cellular subsets were sorted and processed for mini-RNAseq. Inasmuch as at week-1 after grafting LMDPs expressed homogeneously high CD117 levels, they were not further fractionated (unf: unfractionated); due to limited cell yields week-2 LMDPs were divided into CD117^hi^ or CD117^lo^ subfractions. (B). Curves show the dynamic of 20 selected genes across all time points and cellular compartments; gene expression levels are smoothened, normalized, and scaled.

Taken as a whole, these data indicate that increasing expression of B-lineage genes (detectable as early as the level of HSCs) drives the temporal shift of lymphoid potential toward the CD127^+^ ELP subset.

### CD117^lo^ MLPs follow independent stepwise or direct differentiation pathways

To characterize the route of CD117^lo^ MLP differentiation, we next followed the changes in lymphoid potential and clonal architecture taking place across 5 consecutive compartments defined along a continuum of CD45RA and CD117 expression (Fig. 6A and Extended Data Fig. 1A). Corresponding CD34^hi^ HSPC fractions were thus sorted from HSC-engrafted mice at week-3 and cultured as above under the clonal diversification condition. Analysis of 1,853 clones derived from 2,410 cells showed the expected decrease of median clone size as cells progress along the CD45RA/CD117 lymphoid axis and confirmed that ELP production patterns are largely independent of the ancestor cell type (Extended Data Fig. 8A-C). Applying the UMAP algorithm to this compendium showed that HSCs mostly comprise multipotent progenitors whereas downstream CD45RA^int^ MPPs or CD117^hi-int^ LMDPs display a gradual enrichment in lymphoid progenitors whose percentages reach maximum levels in CD117^lo^ MLPs (Fig. 6B and Extended Data Fig. 8D-F). Stratification of clones according to lineage output confirmed that multipotent and granulomonocytic progenitors distribute across the first three compartments. Also consistent with previous observations, a minor subset of HSCs with exclusive self-renewal activity was detected within the CD45RA^-^Lin^-^ compartment. Clone stratification confirmed that amongst LMDPs decreasing expression of CD117 marks the loss of granulomonocytic potential, consistent with previously analyses performed at the population level. CD117^int^ LMDPs comprised a majority of bi- or unipotent lymphoid or dendritic progenitors, reinforcing the idea that the separation between lymphoid and dendritic lineage fates preferentially takes place at this level. Again, CD117^lo^ LMDPs exclusively comprised lymphoid-restricted progenitors. Hierarchical clustering of the clone compendia according to differentiation potentials further confirmed the developmental promiscuity between lymphoid and dendritic lineage fates (Fig. 6C and Extended Data Fig. 8G). Collectively, these results indicate that lymphoid specification follows a stepwise hierarchy characterized by early segregation of granulomonocytic lineage fates, followed by individualization of lympho-dendritic progenitors and then by downstream segregation into unipotent dendritic or lymphoid progenitors

**Figure 6.**
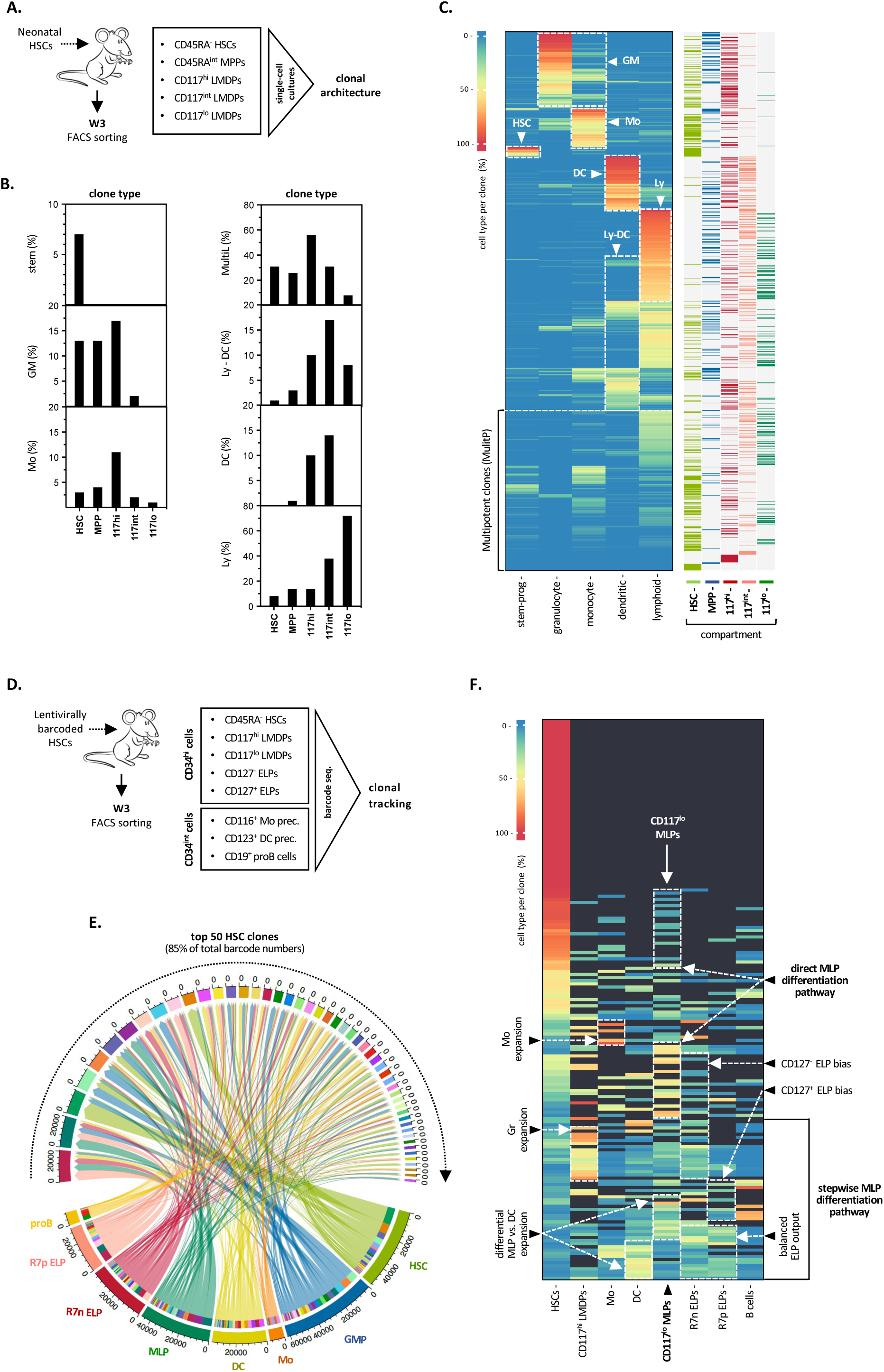
Multimodal mapping of MLP differentiation pathways. (A-D). In vitro assessment of the clonal architecture of HSC, MPP, CD117^hi-int^ LMDP and CD117^lo^ MLP compartments defined along a continuum of CD45RA and CD117 expression. (A) Experimental design: the cell subsets were sorted from HSC-xenografted mice at week-3 and cultured for 14 days under clonal conditions before quantification of lineage output; results are pooled from a representative experiment. (B) Bar plots show the percentages of stem (CD34^+^Lin-), granulomonocytic (GM), monocytic (M), multi-lineage (MultiL), lympho-dendritic (Ly-DC), dendritic (DC) and lymphoid (Ly) clones within the indicated cellular fractions (see also legend of Figure 4). (C) Heatmap shows the lineage output of a compendium of 1853 clones derived from the indicated cell subsets; percentages of the different cell types are normalized relative to total hu-CD45^+^ cells; percentages of lymphoid cells are defined as the sum of CD19^+^ BL and CD127^-/+^ ELP percentages; hierarchical clustering uses Euclidian distance; right panel show the projection of individual clones derived from each population. Dashed areas indicate lineage and cell type-specific production patterns (HSC: self-renewing HSCs; MPP: multipotent lympho-myeloid; GM: granulomonocytic; Mo-DC: mono-dendritic; DC-Ly: dendritic cell-lymphoid; DC: dendritic cell; Ly: lymphoid). (D-F). Lineage output of barcode-labeled HSCs transplanted to NSG mice: (D) Experimental design: neonatal HSCs isolated were transduced with a library of barcoded gfp-reporter lentiviruses (transduction efficiency < 30%) before intravenous injection to irradiated NSG mice (2 x 10^5^ cells/mouse); recipient mice were sacrificed as above at week-3 after grafting; the indicated hu-CD45^+^egfp^+^ subsets were FACS-sorted and processed for analysis of barcode repertoires. (E) Circos plot shows the lineage output of the top 50 largest HSC clones representing 85% of total barcode numbers. (G) Heatmap representation of the lineage output of 179 active HSC clones; clustering uses complete linkage and Euclidian distance; dashed areas show lineage-specific production patterns. Gr: granulocyte; Mo: monocyte; DC: dendritic cell; Ly: lymphoid; MLP: multi-lymphoid.

A lentiviral barcoding approach was next applied to trace the origin of CD117^lo^ MLPs (Fig. 6D). Analysis of the barcode repertoire distributed across eight cellular subsets isolated from mice reconstituted with barcoded HSCs allowed detection of 1,490 unique barcodes from which only 179 (12%) were present within the HSC compartment indicating that the majority of barcoded HSCs initially engrafted have been lost (Table S9). Consistent with this, 30% of remaining barcoded HSCs lacked detectable progeny suggesting functional quiescence. The 50 largest HSC clones accounted for 85% of the total barcodes, an indication that 3 weeks after grafting only 3% of initially engrafted HSCs contributed to ongoing hematopoiesis (Fig. 6E). Consistent with the functional data, analysis of barcode frequencies across the progeny of active HSCs (Fig. 6F) showed that expansion of the granulomonocytic compartment coincides with decreased dendritic and lymphoid outputs and confirmed that the barcode numbers detected in dendritic precursors and CD117^lo^ MLPs vary in inverse proportions which, once again, is suggestive of differential regulation between lymphoid and dendritic fates at the level of common upstream precursors. As expected, these analyses also confirmed that 3 weeks after grafting CD117^lo^ MLPs comprise a mixture of clones with balanced or biased ELP outputs. To our surprise, analysis of barcode repertoires also revealed that a significant fraction of CD117^lo^ MLPs originate from HSCs with barely detectable myeloid or dendritic progeny which was indicative of a direct differentiation.

Taken together, these data show that CD117^lo^ MLPs follow independent stepwise or direct differentiation pathways. Also, they indicate that the routes of lymphoid differentiation are not constrained by the hematopoietic hierarchy.

### Single-cell transcriptional mapping of CD117^lo^ MLP differentiation pathways

To further investigate CD117^lo^ MLP differentiation and search for pathway-specific gene signatures, CD34^hi^ HSPCs were subdivided into four compartments including the CD127^-^ and CD127^+^ ELPs before analysis by single-cell RNA sequencing (Extended Data Fig. 1C). This procedure ensures an equal representation of the corresponding cell fractions, thereby maximizing the capture of discrete cellular states. After quality control and doublet exclusion the final dataset was composed of 37,092 cells which were projected on a 2D space (UMAP) (Fig. 7A and Extended data Fig. 9A).

**Figure 7.**
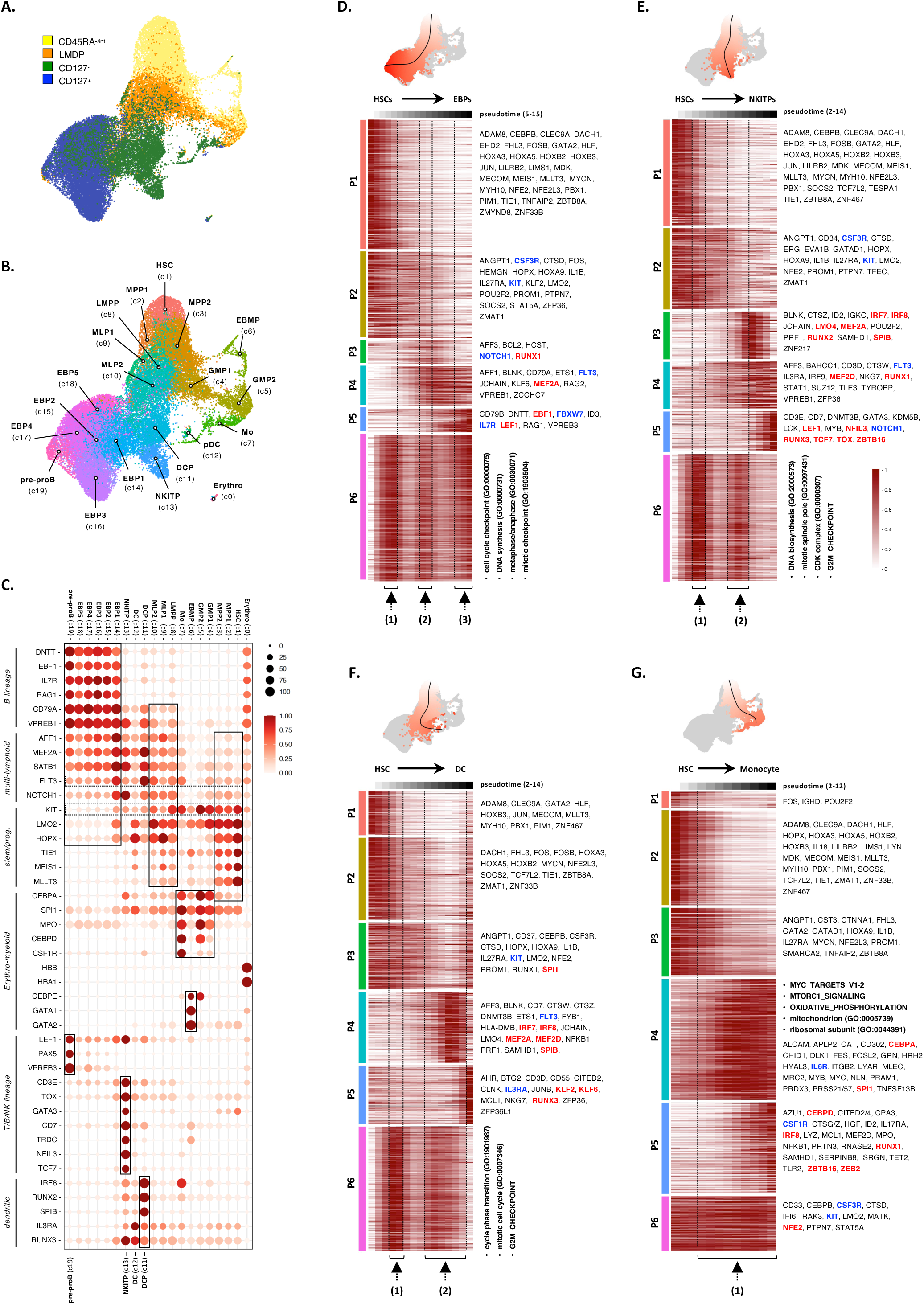
In silico reconstruction of lymphoid differentiation trajectories. The indicated CD45RA^-/int^, CD45RA^hi^ and CD127^-^ or CD127^+^ HSPC subsets were isolated from the BM of HSC-xenografted mice (week-3) and subjected to scRNA-seq profiling with the 10X genomic platform. (A, B). UMAP visualization of the compendium of CD34^+^ HSPCs colored by (A) cellular compartment or (B) cell clusters (c0-19) annotated based on expression of canonical lineage marker genes for hematopoietic stem cells (HSC: C1), multipotent progenitors (MPP: C2-3), granulomonocytic progenitors (GMP: C4-5), eosinophil/basophil/mast cell progenitors (EBMP: C6), monocytes (Mo: C7), lympho-primed multipotent progenitor (LMPP: C8), multi-lymphoid progenitors (MLP: C9-10), dendritic cell precursors (DCP: C11), plasmacytoid dendritic cells (pDC: C12), NK/ILC/T progenitors (NKITP: C13), early B cell precursors (EBP: C14-18) or pre-proB cells (pre-proB: C19), erythroid progenitors (Erythro: C0). (C). Spot plot based on expression of 42 selected genes show lineage fingerprints of clusters C1-19; dashed areas show cluster-specific gene signatures. (D-G). Variations along pseudotime of lymphoid (D) B or (E) T/NK/ILC or (F) myeloid dendritic or (G) monocyte lineage-affiliated transcriptional programs. Upper UMAPs show lineage-specific differentiation trajectories represented as smoothed black curves; lineage-specific cells are color-coded (white to red) according to pseudotime values. Heatmaps represent min to max normalized expression of dynamically variable genes; trajectory inference analyses were performed with SlingShot. Growth factor receptors are in blue; TFs are in red; GO/GSEA enrichments are indicated. Dashed lines and black arrows correspond to proliferation phases (1-3).

Graph-based Louvain clustering identified 20 clusters (C0-19) (Fig. 7B and Table S10) associated with 13 cell types identified based on a set of 42 manually selected lineage gene markers among the 3,844 DEGs (Fig. 7C). Analysis of cluster-specific signatures confirmed the overexpression of genes involved in stem cell maintenance or self-renewal (*LMO2*, *HOPX*, *TIE1*, *MEIS1*, *MLLT3*) in HSCs (C1) and the expected enrichment in protein synthesis or oxidative phosphorylation pathways in downstream multipotent progenitors (MPP1/2; C2-3). The MPPs that also displayed evidence of onset of lymphoid (MPP1 [C2]: *AFF1*, *SATB1*, *FLT3*) or myeloid (MPP2 [C3]: *CEBPA*) transcriptional priming (Fig. 7C). Granulomonocytic progenitors (GMP1-2; C4-5) overexpressed *KIT* transcripts and granulocytic lineage genes (*CEBPA*, *CSF3R*, *MPO*, *ELANE*), while subsequent individualization of lineage-specified monocyte precursors (C7) was associated with the upregulation of *CSF1R*, *CEBPD*, *IRF8* or *SAMHD1*. Eosinophil/Basophil/Mast cell progenitors (EBMP; C6: *GATA1*, *GATA2*, *CEBPE*, *IL5RA*) remained confined to the CD45RA^-/int^ compartment indicating that their independent differentiation (Extended data Fig. 9A) ^36^. Consistent with the mini-RNA-seq data, enrichment analyses confirmed that upregulation of pre-BCR components in actively cycling LMPPs (C8). Most importantly, these analyses also allowed the identification of distinct MLP1 (C9) and MLP2 (C10) clusters with distant projection on the transcriptional space, but largely overlapping G1 cell-cycle statuses and gene signatures (*AFF1*, *MEF2A*, *SATB1*, *FLT3*, *CD79A*, *VPREB1*) (see also Extended data Fig. 9B). Interestingly, MLP1s displayed higher expression of stem cell-associated transcripts (*HOPX*, *TIE1*, *MEIS1*, *MLLT3*) which strongly suggests that they arise from the direct MLP differentiation pathway. Consistent with this idea, an accumulation of cells expressing T/NK/ILC (*NFIL3*, NKG7, *CD7, CCR7*) or B (*EBF1*, *CD79A*, *IGHD*, *JCHAIN*) lineage markers was detected in the vicinity of MLP1s (C9) (Extended data Fig. 9C). In contrast, the spatial proximity of MLP2s (C10) with LMPPs (C8) strongly suggests that they follow the stepwise differentiation pathway.

Louvain clustering subdivided downstream CD127^+^ ELPs into immature early B-cell progenitors (EBP1; C14) displaying a dual multi-lymphoid (*FLT3*, *SATB1*, *AFF1*, *MEF2A*) and B-lineage (*IL7R*, *EBF1*, *SOX4*, *ZCCHC7*, *RAG2*, *DNTT*) signature, and four prototypic EBP2-5 subsets (C15-18) converging toward the pre-proB stage (C19: *LEF1*, *PAX5*, *VPREB3*). As expected ^31^, NK/ILC/T progenitors (NKITP; C13) overexpressed NK-ILC (*ID2*, *NFIL3*, *TOX*, *LMO4*) or T (*LEF1*, *TCF7*, *GATA3*, *TRDC*, *CD3E*) lineage genes and they were exclusively situated within the CD127^-^ compartment. The finding that dendritic cell progenitors (DCP) (C11: *IRF8*, *RUNX2*, *SPIB*, *FLT3*, *IL3R*) retained expression of B-lymphoid genes (*IGKC*, *BLNK*, *IGHM*, *JCHAIN*) stresses their developmental promiscuity with the lymphoid lineage. Notably, LMPPs (C8) comprised a minor contingent of *IRF8^+^* cells, reinforcing the view that segregation of the lymphoid and dendritic potentials occurs at this level (Extended Data Fig.9D, left panels).

Analysis of the expression pattern of 20 lineage marker genes in individual cells confirmed that differential commitment toward the granulomonocytic or lymphoid lineages is associated with extinction of heterologous differentiation programs (Extended Data Fig.10). Consistent with discrepant cell-intrinsic versus -extrinsic regulation of underlying regulatory networks, these analyses revealed that, whereas sterile *IGHM*/*D* transcripts are detected as early as the HSC level, the expression of *TRD, NFIL3*, *TCF7* or *GATA3* remains confined to NKITPs (C13).

At last, detection of distinct MLP1 and MLP2 clusters by unsupervised Graph-based Louvain clustering confirms that CD117^lo^ MLPs arise from independent direct or stepwise differentiation pathways.

### Lymphoid trajectories comprise alternating phases of cell differentiation and proliferation

To gain an overview of lymphoid differentiation trajectories, we finally performed lineage reconstructions with pseudotime inferences using the SlingShot method ^37^ on our compendium of 37,092 cells (Table S11).

These analyses revealed that lymphoid differentiation starts with the sequential downmodulation of stem/progenitor (P1) and erythro-myeloid lineage (P2) genes followed by the proliferative burst of LMPPs (Fig. 7D, E). Also, they confirmed that a proliferation arrest (P6) occurring at the level of CD117^lo^ MLPs (C10) precedes the dichotomic choice between B and NK/ILC/T lymphoid lineages. Further dissection of the B-cell pathway identified three developmental intermediates sequentially upregulating B-lineage TFs and marker genes (P3-5) and revealed that B-cell progenitors undergo two rounds of proliferation (P6). The first expansion wave immediately downstream of MLPs drives progression towards EBP1s whose proliferation arrest coincides with *NOTCH1* and *FLT3* repression and corresponding upregulation of *IL7R* and *FBXW7* transcripts. After a second round of proliferation presumably driven by IL7, B-cell progenitors reach the pre-proB stage and start to rearrange the IGH locus. Differentiation along the NK/ILC/T pathway initiates with promiscuous expression of lymphoid or dendritic lineage genes (P3-4) followed by the upregulation of TFs controlling NK/ILC/T lineage fates (P5). In contrast to its B-lineage counterpart, the NK/ILC/T differentiation pathway comprised of a single wave of cell proliferation (P6) driven by Flt3 signaling. Subsequent analysis of the dendritic pathway confirmed developmental promiscuity with the lymphoid lineage and found that upregulation of *IL3R* precedes the segregation into plasmacytoid and conventional DC subsets (Fig. 7F and Extended Data Fig. 9D, right panels). At last, single-cell regulatory networks inferences showed that myeloid monocytes and granulocytes follow essentially continuous differentiation pathways driven by gene differentiation and proliferation modules operating in a combinatorial manner (Fig. 7G and Extended Data Fig. 11A), findings consistent with an earlier report ^38^.

Collectively, these results show that lymphoid differentiation pathways are characterized by alternating phases of proliferation and differentiation. The finding that cell-proliferation arrests coincide with transitions in differentiation states and growth factor dependencies suggests that they might correspond to previously undescribed lymphoid development checkpoints.

## DISCUSSION

In this study, we establish a multimodal map of human lymphopoiesis at clonal resolution that extends the current view of lymphoid organization (Extended data Fig. 11B).

Our results demonstrate that previously characterized CD127^-^ and CD127^+^ ELPs ^31^ differentiate from a novel subset of CD117^lo^ MLPs. We also provide evidence that the CD117^lo^ MLPs arise from independent stepwise or direct differentiation pathways. Whereas the direct MLP differentiation pathway proceeds in absence of multilineage diversification, its stepwise counterpart follows the hematopoietic hierarchy and is characterized by an early separation of granulomonocytic potential, followed by individualization of bipotent lympho-dendritic progenitors and then by downstream segregation into unipotent dendritic or lymphoid progenitors. Although it currently remains unclear whether these two MLP differentiation pathways are redundant and if the corresponding cell subsets display common or pathway-specific functional properties, our data indicate that the route of lymphoid specification is not constrained by the hematopoietic hierarchy.

Transcriptional mapping of lymphoid specification performed at both the population and single-cell levels revealed that, whereas divergence with the myeloid lineage is observed immediately downstream of HSCs, lymphoid priming starts at the level of actively proliferating CD117^int^ LMDPs which correspond to the human counterparts of mouse LMPPs ^39^. Gene expression analyses confirmed that lymphoid commitment of downstream CD117^lo^ MLPs is associated with repression of myeloid lineage genes coinciding with upregulation of genes coding for early lymphoid regulators and pre-BCR components. These analyses also revealed that CD117^lo^ MLPs retain the expression of stem cell-associated genes which might reflect an imprint left by their upstream precursors (particularly when they arise from the direct differentiation pathway) or indicate that lymphoid restriction is associated with the acquisition of stem cell-like properties. The finding that CD117^lo^ MLPs also decrease expression of cell cycle-related genes is consistent with our observation that they expand poorly in diversification assays irrespective of the growth factor combination tested. These data are concordant with earlier reports showing that common or multi-lymphoid progenitors display limited in vitro and in vivo amplification ^20, 23^. Since our results demonstrate that a proliferation arrest precedes the dichotomic choice between CD127^-^ (NK/ILC/T) and CD127^+^ (B) lymphoid pathways, we propose that the CD117^lo^ MLP stage marks the first lymphoid development checkpoint described so far. Notably, CD117^lo^ MLPs might also represent a critical point of vulnerability for leukemogenic transformation ^40^. From an evolutionary perspective and based on shared immunophenotypes, gene signatures and functional properties, CD117^lo^ MLPs could be viewed as human counterparts of mouse CD117^lo^FLT3^hi^ lymphoid-primed progenitors (LPPs) ^17^. Given the evidence that mouse early T-lineage progenitors (ETPs) differentiate independently of CD127^+^ CLPs ^41–43^, it is possible that the bifurcation between NK/ILC/T- and B-lymphoid lineage in mice also takes place at the level of LPPs. If so, this would also imply that the bipartite organization of lymphoid architecture ^31^ is conserved across species.

Analysis of the developmental relationships between CD127^-^ and CD127^+^ ELPs led to the unexpected finding that they are subjected to a differential cell-extrinsic versus -intrinsic regulation. Indeed, whereas Flt3 signaling drives CD117^lo^ MLPs toward the NK/ILC/T differentiation pathway in an instructional manner, the emergence of B lineage-biased CD127^+^ ELPs depends exclusively on cell-intrinsic regulatory mechanisms. The proposition that entry into the B-lymphoid pathway is regulated cell-autonomously relies on several lines of evidence. Firstly, our data indicate that the time-dependent increase in CD127^+^ ELP production observed in xenografted mice is driven by a global B-lineage shift in the lymphoid potential of their precursors, which is also observed as an increase in expression of B-lymphoid genes that can be detected as early as the HSC level. Secondly, they provide evidence that these changes are ultimately conditioned by the divisional history of CD34^+^ HSPCs, independent of their differentiation stage. Third, a functional screen of lymphoid regulators failed to identify soluble inducers of CD127^+^ ELP differentiation. Functional characterization of CD117^lo^ MLPs subsequently revealed that their intrinsic B-lineage bias is counterbalanced by an exquisite susceptibility to Flt3L. Thus, consistent with earlier reports that Flt3L is a key regulator of lymphoid development ^16, 44, 45^, our results indicate that Flt3 signaling finely tunes the NK/ILC/T- versus B-lineage choice at the level of CD117^lo^ MLPs. At present it remains unclear whether and how these fundamentally distinct modes of regulation influence the pattern of lymphocyte production across development and aging (see also by Keita et al., submitted).

Finally, reconstruction of lineage trajectories confirmed that, in contrast to the granulomonocytic differentiation pathway that proceeds in a continuous manner ^38, 46^, lymphoid differentiation trajectories are intrinsically discontinuous and comprise alternating phases of cell proliferation and differentiation that amplify the pools of lymphoid progenitors prior to the onset of antigen receptor rearrangement.

## AUTHOR CONTRIBUTIONS

K. A.H. designed the study and wrote the paper; E. C. & S. L. performed experiments; G. P A. & S.D. analyzed the clonal and fate mapping data; E. N., E. V., G. P. A. & K. C. conducted the fate mapping studies and barcoding analyses; F. C. performed the transcriptome analyses and wrote the paper; E.A.M. & Z. K. analyzed the data and wrote the paper; B. C. ensured the scientific supervision of the project.

## ACKNOWLEDGEMENTS

The authors thank Christelle Doliger and Niclas Setterblad (Plateforme d’Imagerie et de Tri Cellulaire, IRSL, Paris France). We are grateful to Laurent David (Inserm 1064, Nantes) and Bernard Jost (GenomEast platform, IGBMC, Strasbourg, France). We thank Justine Poirot for help in single cell analyses. We also thank Sophie Ezine and Bela Papp for critical discussions. This work was supported by the Agence de la Biomédecine, Agence Nationale de la Recherche (ANR EpiDev), the Institut National du Cancer (InCA B-REC) and by the INSERM HuDeCA network.

The authors have no conflict of interest.

## SUPPLEMENTARY TABLES

**Table S1**. Gene expression profiling of 4 HSPC subsets isolated from mice at week 3 after grafting. (A) Distribution of 207 differentially expressed genes (DEGs) across 14 clusters (C1-14; fold-change ≥ 1.5; adjusted F-value ≤ 0.005); population-specific gene set enrichments are highlighted in gray. (B) List of GO and GSEA categories enriched in clusters P1-14 (adjusted *p* values are indicated).

**Table S2**. List of (A) Taqman probes and (B) soluble ligands used in this study.

**Table S3**. Transcriptional response of 4 HSPC subsets to short-term stimulation with Flt3L. (A) Distribution of 638 DEGs across 9 clusters (P1-9; fold-change ≥ 1.5; adjusted F-value ≤ 0.005); population-specific gene set enrichments are highlighted in gray. (B) List of GO and GSEA categories enriched in Flt3-treated versus untreated HSPCs (adjusted *p* values are indicated).

**Table S4**. Transcriptional response of 4 HSPC subsets to short-term stimulation with TPO. (A) Distribution of 544 DEGs across 4 clusters (Q1-5; fold-change ≥ 1.5; adjusted F-value ≤ 0.005); population-specific gene set enrichments are highlighted in gray. (B) List of GO and GSEA categories enriched in in TPO-treated versus untreated HSPCs (adjusted *p* values are indicated).

**Table S5**. Temporal variations of the gene expression profile of HSCs isolated from mice between week 1-5 after grafting (P1-5; 480 DEGs). (A) Distribution of 480 DEGs across 5 clusters (P1-5; fold-change ≥ 1.5; adjusted F-value ≤ 0.005); (B) List of GO and GSEA categories enriched in clusters P1-5 (adjusted *p* values are indicated).

**Table S6**. Temporal variations of the gene expression profile of CD117^hi^ LMDPs isolated from mice between week 1-5 after grafting (P1-5; 501 DEGs). (A) Distribution of 501 DEGs across 5 clusters (P1-5; fold-change ≥ 1.5; adjusted F-value ≤ 0.005); (B) List of GO and GSEA categories enriched in clusters P1-5 (adjusted *p* values are indicated).

**Table S7**. Temporal variations of the gene expression profile of CD117^int^ LMDPs (also referred to as LMPPs) isolated from mice between week 1-5 after grafting (P1-5; 612 DEGs). (A) Distribution of 612 DEGs across 5 clusters (P1-5; fold-change ≥ 1.5; adjusted F-value ≤ 0.005); (B) List of GO and GSEA categories enriched in clusters P1-5 (adjusted *p* values are indicated).

**Table S8**. Temporal variations of the gene expression profile of CD117^lo^ LMDPs (also referred to as MLPs) isolated from mice between week 1-5 after grafting (P1-5; 871 DEGs). (A) Distribution of 871 DEGs across 5 clusters (P1-5; fold-change ≥ 1.5; adjusted F-value ≤ 0.005); (B) List of GO and GSEA categories enriched in clusters P1-5 (adjusted *p* values are indicated).

**Table S9.** Distribution of barcode copy numbers across eight cellular subsets isolated at week-3 after grafting from mice reconstituted with HSCs transduced with a lentiviral barcode library.

**Table S10.** Single-cell transcriptional profiling of 37,092 HSPCs isolated from mice at week-3 after grafting. Distribution of 3,844 differentially expressed genes (DEGs) across 20 clusters identified by graph-based Louvain clustering (C1-20).

**Table S11.** Single-cell transcriptional mapping of 5 lympho-myeloid differentiation trajectories. (A) Lineage-specific pseudotime Slingshot clusters (P1-6); (B) List of the top 30 GO and GSEA categories enriched in Lineage-specific Slingshot clusters (adjusted *p* values are indicated).

## METHODS

### Human Sample Collection

Umbilical cord blood (UCB) was provided by the Unité de Thérapie Cellulaire of Hôpital Saint-Louis (Paris).

### Mice

NOD.Cg-Prkdc*^scid^*IL2RG*^tm1wjl^*/SzJ (005557) mice known as NOD scid gamma (NSG) mice (Jackson Laboratory, Bar Harbor, MI) were housed in the pathogen-free animal facility of Institut de Recherche Saint Louis (Paris). Female NSG mice were xenografted at 2 months of age. The Ethical Committee at Paris Nord University approved all performed experiments.

### Processing of human cord blood and cell separation

Blood cells were separated by Ficoll-Hypaque centrifugation (Pancoll, PAN Biotech GmbH) before processing for flow cytometry, cell sorting or CD34^+^ HSPC isolation. Human CD34^+^ cells were isolated with the CD34 Microbead kit (Miltenyi Biotech; purity >90%), frozen in heat-inactivated fetal calf serum (FCS) supplemented with 10% DMSO and stored in liquid nitrogen until use.

### Xeno-transplantations

Xeno-transplantations were performed essentially as described ^31^. NOD scid gamma (NSG) mice (Jackson Laboratory, Bar Harbor, MI) housed in the pathogen-free animal facility of Institut de Recherche Saint Louis (Paris) were irradiated (2.25 Gy) 24-hr before injection of neonatal CD34^+^ cells in the caudal vein. Recipient NSG mice were sacrificed between week-1 and week-5 after grafting. The Ethical Committee at Paris Nord University approved all performed experiments.

### Flow cytometry and cell sorting

Single-cell suspensions recovered from the femurs and tibias of xenografted mice were filtered through a cell strainer (70 µm; BD biosciences) and depleted in mouse cells using the Mouse Cell Depletion Kit (Miltenyi) before being processed for Flow Cytometry. Human single-cell suspensions were incubated with human Fc receptor-binding inhibitor (Fc Block, eBioscience) before surface staining with anti-human monoclonal antibodies (mAbs). Fluorescence minus one (FMO) controls with control isotype antibodies were used to define positive signals for flow cytometry or cell sorting. Dead cells were excluded with the Zombie Violet Fixable Viability Kit (Biolegend). For labeling, cells were resuspended in PBS, 2 % FCS (1 to 5 x10^7^ cells/500 μl) and incubated with the following mAbs: CD45 AF700 (Biolegend, clone HI30), CD34 PB (Biolegend, clone 581), CD7 FITC (Beckman Coulter, clone 8H8.1), CD38 PerCPCy5.5 (Biolegend, clone HIT2), CD123 BV785 (BD Bioscience, clone 7G3), CD115 APC (Biolegend, clone 9-4D2-1E4), CD45RA PE (BD Bioscience, clone HIT100), ITGB7 PC7 (eBioscience, clone FIB504), CD33 PE-CF594 (BD Bioscience, clone WM53), CD19 BV711 (BD bioscience, clone HIB19), CD116 APC-vio770 (Miltenyi, clone REA211), CD127 PC5 (Biolegend, clone A019D5), CD117 BV605 (Biolegend, clone 104D2), CD10 BV650 (BD bioscience, clone HI10a), also CD7 PE-CF594 (BD Bioscience, clone M-T701). Flow cytometry and cell sorting were performed with a BD Fortessa Analyzer or a BD FACSAria III sorter (BD Biosciences; (purity ≥ 95%). Flow cytometry analyses were performed using the FlowJo software (Version 10.7).

For investigating the effect of conservative cell division on the lymphoid differentiation potential, neonatal HSC or LMDPs sorted from the UCB were labeled with proliferation-dependent dye carboxyfluorescein diacetate succinimidyl ester (CFSE) using the Vybrant® CFDA SE Cell Tracer Kit (Life Technologies) prior injection to irradiated NSG mice (2 x 10^5^ cells/mouse). Mice engrafted with the CFSE-labeled cells were sacrificed 8 day later before huCD45^+^ BM cells were labelled with the same antibody cocktail as above except for the CD7 antibody (CD7 PE-CF594; BD Bioscience, clone M-T701).

### Cell cultures and analysis

For multi-lineage diversification assays under bulk conditions (100 cell/well), OP9 or OP9-DL4 stromal cells were seeded in Opti-MEM-Glutamax supplemented with 2.5% FCS, 7.5% BIT 9500 (StemCell Technologies), 1% penicillin-streptomycin and 1/1000 *β*-mercapto-ethanol (Life Technologies) in 96-well U-bottom plates 24 hours prior to co-culture with the indicated combinations of growth/differentiation factors (10 ng/ml each; all from Miltenyi Biotec). CD34^+^ HSPCs isolated from the BM of xenografted mice sacrificed between weeks 1-5 after grafting were directly seeded by FACS in the 96-well plates and cultured for 7 or 14 days before quantification of lineage outputs. Clonal diversification assays were conducted for 14 days under the same condition, except that cultures were supplemented with SCF, TPO and GM-CSF (10 ng/ml each).

For assessment of lineage output, in vitro-differentiated cells were labeled with the following antibodies: CD45 AF700 (Biolegend, clone HI30), CD34 PB (Biolegend, clone 581), CD7 FITC (Beckman Coulter, clone 8H8.1), CD123 BV786 (BD Bioscience, clone 7G3), CD115 APC (Biolegend, clone 9-4D2-1E4), CD45RA PE (BD Bioscience, clone HIT100), ITGB7 PC7 (eBioscience, clone FIB504), CD24 PE-CF594 (BD Bioscience, clone ML5), CD19 BV711 (BD bioscience, clone HIB19), CD116 APC-vio770 (Miltenyi, clone REA211), CD127 PC5 (Biolegend, clone A019D5), CD15 BV605 (BD Bioscience, clone HI10a), CD10 BV650 (BD bioscience, clone HI98). Flow cytometry analyses were performed with a BD Fortessa Analyzer and the FlowJo software.

### Transcriptional profiling by multiplex PCR

Gene expression analyses were performed with the Fluidigm 96.96 Dynamic Array IFC and TaqMan Gene Expression Assays (Life Technologies). The indicated cell subsets were sorted by 50 cell pools directly into 96-well PCR plate containing 2.5 ml TaqMan specific gene assay mix (Applied Biosystems), 5 ml of CellsDirect 2x Reaction mix, 0,2 ml SuperScript TM III RT/PlatinumR Taq Mix (Invitrogen, CellsDirect one-step qRT-PCR kit), 1.2 mL TE buffer, and 0.1 ml SUPERase-In RNase Inhibitor (Ambion). Reverse transcription was performed for 15 min at 50°C followed by 2 min at 95°C for RT inactivation before cDNAs were pre-amplified for 21 cycles at 95°C for 15 sec and 60°C for 4 min. Preamplified products were diluted 1:5 in TE buffer and analysed on a Biomark system (Fluidigm) with the following PCR cycling condition: 95°C for 10 min and 40 cycles at 95°C for 15 sec and 60°C for 60 sec. Data were analysed using the Biomark qPCR Analysis software (Fluidigm). For quantification of gene expression, data were analyzed by the DDCt method. Results were normalized relative to HPRT expression and expressed as mean expression level of ≥ 6 biological replicates. Hierarchical clustering was performed on standardized means of gene expression levels based on Euclidean distance using the ‘‘pheatmap’’ R package. A complete list of the TaqMan primers is provided as Table S2.

### Gene expression profiling by mini-RNA-seq

The protocol used for mini-RNAseq was adapted from the method developed by Soumillon et al. ^47^. For each cell population batches of 100 cells (2-10 replicates) were sorted by FACS in 96-well V-bottom plate wells containing 2 μl of lysis buffer (Ultra-pure Water, 10% Triton X-100, RNasinPlus 40U/ul; Promega). After evaporation of the lysis buffer at 95°C for 3 min, first strand cDNA synthesis was performed using the Maxima H Minus Reverse Transcriptase (Thermo Fisher), E3V6NEXT primers (specific primer for each well) and Template Switching Primer: E5V6NEXT. Resulting cDNAs were pooled and purified by the DNA Clean and Concentrator TM-5 kit (Zymo research) and treated with Exonuclease I (New England Biolabs) before amplification using the Advantage 2 PCR kit (Clontech) and the SINGV6 primer (95°C for 1 min, 15 cycles at 95°C for 15 sec, 65°C for 30 sec, 68°C for 6 min, and 72°C for 10 min). PCR products were purified by Agencourt AMPure XP (Beckman Coulter). Libraries were prepared using the Illumina DNA Prep kit (Illumina) according to the manufacturer’s guidelines and sequenced on a HiSeq 4000 (Illumina) at the Genome East Platform of the IGBMC (Strasbourg, France). Image analysis and base calling were performed using RTA 2.7.7 and bcl2fastq 2.17.1.14 software. Adapter dimer reads were removed using Dimer Remover (https://sourceforge.net/projects/dimerremover/).

### Single cell-RNA-seq

For the droplet encapsulation 2,5 x 10^4^ live, single, HSC/MPP (CD45RA^-/int^), LMDP (CD45RA^hi^ITGB7^-^), CD127^-^ (ITGB7^lo/+^) or CD127^+^ (ITGB7^-/+^) cells isolated by FACS from the BM of NSG-UCB mice at week-3 after grafting were loaded onto each channel of a Chromium chip before encapsulation on the Chromium Controller (10x Genomics). Single-cell sequencing libraries were generated using the Single Cell 3’ Kit v3.1 (10X Genomics) according to the manufacturer’s protocol. Sample quality was controlled with a Bioanalyzer Agilent 2100 using a High Sensitivity DNA chip (Agilent Genomics). Libraries were sequenced using an Illumina HiSeq 4000 to achieve a minimum depth of 100,000 raw reads per cell.

### Lentiviral barcoding experiment

The barcoded GFP-BC32 lentiviral-vector library was kindly provided by K. Cornils (University Medical Center, Hamburg, Germany) ^48^. UCB CD34^+^ cells (purity > 90%) were transduced 6-8 hours with the barcoded lentiviruses used at a multiplicity of infection allowing approximately 30% of transduction efficiency to ensure that that each cell contains only one integrated barcode. Infections were performed in RPMI medium supplemented with 20% BIT 9500 (StemCell Technologies) and SCF (50 ng/ml), FLT3L (50 ng/ml), TPO (20 ng/ml) and IL-3 (10 ng/ml; all from Miltenyi). NSG mice reconstituted with the lentivirally-transduced CD34^+^ cells were sacrificed at week-3 after grafting before harvesting of BM and sorting of hu-CD45^+^ cellular subsets, as above. To investigate the barcode content, the BC32 sequences were subjected to two rounds of amplification with common (DUAL_P5_01) or population-specific (MLPX35-42) primers. Barcode libraries were then pooled and sequenced with Illumina HiSeq 4000. Barcode analyses were performed as described with the genBaRcode package (https://cran.r-project.org/web/packages/genBaRcode) ^49^.

### Quantification and statistical analyses

#### Algorithm-assisted classification of in vitro-derived clones

Clonal data were processed with the ClonAn software developed by S. Diop. Analyses were based on the UMAP package ^50^.

#### Mini-RNA-seq and bioinformatics

Data quality control and pre-processing were performed by SciLicium (Rennes, France). Briefly, the first read contains 16 bases that must have a quality score higher than 10. The first 6 bp correspond to a unique sample-specific barcode and the following 10 bp to a unique molecular identifier (UMI). The second reads were aligned to the human GRCh38/hg38 reference genome from the UCSC website using BWA version 0.7.4.4 with the parameter “−l 24”. Reads mapping to several positions in the genome were filtered out from the analysis. The complete pipeline has been previously described in ^51^. After quality control and data pre-processing, a gene count matrix was generated by counting the number of unique UMIs associated with each gene (lines) for each sample (columns). The UMI matrix was further normalized with the regularized log (rlog) transformation package implemented in the DeSeq2 package ^52^. The statistical comparisons were performed in the AMEN suite of tools ^53^. For each analysis relevant comparisons were selected to identify differentially expressed genes (DEGs). Briefly, genes showing an expression signal higher than 0.0 and at least a 2.0-fold-change between the two experimental conditions of each pairwise comparison were selected. The empirical Bayes moderated t-statistics implemented into the LIMMA package (F-value adjusted using the Benjamini & Hochberg (BH) False Discovery Rate approach, p ≤ 0.05) ^54, 55^ ^56, 57^ was used to define the set of genes showing significant statistical changes across all comparisons of a given transcriptomic analysis. The resulting set of DEGs were then partitioned into distinct expression clusters with the k-means algorithm and presented as heatmaps by using the pheatmap R package developed by R. Kolde (2019) (version 1.0.12. https://CRAN.R-project.org/package=pheatmap). Raw and pre-processed data are accessible at the GEO repository (accession number: GSE215296)

#### Single cell-RNA-seq and bioinformatics

Demultiplexed raw sequencing reads were processed, mapped to the GRCh38 human reference genome and quantified based on unique molecular identifiers (UMIs) with the Cell Ranger pipeline (v3.0.2, 10x Genomics). The resulting raw count matrices were then merged. Subsequently, the Scater R package (v1.10.1) was used to remove outlier cells by using several cell features including the proportions of reads mapping mitochondrial and ribosomal genes. In parallel, doublets were filtered out independently in each individual matrix by using the DoubletFinder R package (v.2.0.2). Cells with less than 1000 detected genes and genes detected in less than 10 cells were removed. Cells from all datasets were assigned a cell cycle phase by using Seurat. Data were normalized using the NormalizeData and the SCTransform functions implemented into Seurat. The top-3000 most varying genes were used to perform a principal component analysis with the RunPCA function implemented in Seurat. Cells were then clustered by using the FindNeighbors and FindClusters functions on the top-30 principal components, with default parameters. Finally, we used the uniform manifold approximation and projection (UMAP) method implemented in Seurat to project single cells in a reduced 2D space. Cell clusters were annotated using a set of 42 known marker genes which are also significantly differentially expressed between cell clusters. Differentially expressed genes were identified with the FindAllMarkers function implemented in Seurat, with default parameters. Cell trajectories were inferred the getLineages followed by the getCurves functions implemented into the SlingShot package ^37^ on the scRNA-seq data by indicating the C1 cell cluster (HSC) as starting cells. Trajectory-based differential expression analysis was performed with the tradeSeq package by using the pseudotime values inferred by SlingShot for each lineage ^37, 58^. Raw and pre-processed data are accessible at the GEO repository (accession number: GSE214950).

---

**Extended Data Fig. 1.**
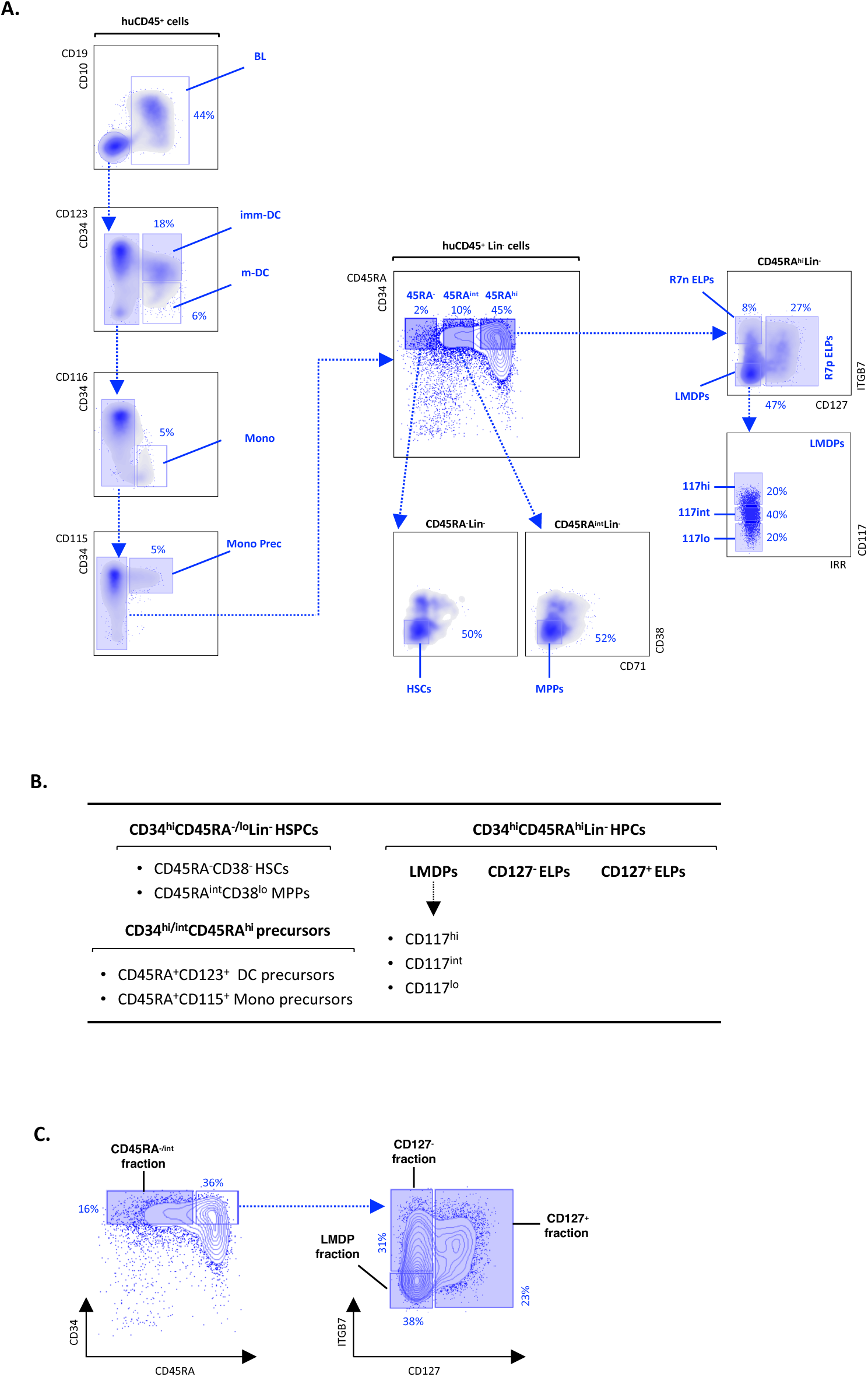
Immunophenotyping profiling and gating procedures. (A). Gating procedure to subdivide hu-CD45^+^ BM cells from xenografted NSG-UCB mice. Bone marrow mononuclear cells (BMNCs) isolated at week 3 HSC-xenografted mice were labeled with our reference 16-antibody panel. After exclusion of doublets gates were set on the human (hu)-CD45^+^ cells (upper left panel). hu-CD45^+^ cell were first partitioned based on CD19 and CD10 before subdivision of the double negative fraction based on CD34 and CD123 expression allowing exclusion of DCs. Remaining CD123^-^ cells were partitioned based on differential CD116/CSF2R expression to exclude monocytes (Mono). After removal of CD34^+^CD115^+^ monocyte precursors, CD34^hi^ HSPCs were partitioned into CD45RA^-^, CD45RA^int^ or CD45RA^hi^ compartments. Within the CD45RA^-^ and CD45RA^int^ compartments, CD38^-^CD71^-^ cell fractions were respectively referred to as the hematopoietic stem cells (HSC) or multipotent progenitors (MPP). CD45RA^hi^ HPCs was first subdivided based on differential ITGB7 and CD127 expression to distinguish Lympho-Myelo-Dendritic Progenitors (LMDPs) from the CD127^-^ (R7n) or CD127^+^ (R7p) Early Lymphoid Progenitors (ELPs). Subdivision of LMDPs based on CD117/c-Kit expression allowed for individualization of three CD117^hi^, CD117^int^ (also referred to as Lymphoid-Primed Mulitpotent Progenitors [LMPPs]) and CD117^lo^ (also referred to as Multi-Lymphoid Progenitors [MLP] fraction) cellular fractions. Bi-dimensional density and dot plots and blue arrows show the gating hierarchy; percentages of the corresponding populations are indicated. Results are representative of > 40 experiments from > 200 xenografted mice. (B). List and nomenclature of the CD34^hi^ HSPCs subsets analyzed in this study. (C). Gating procedure for scRNA-seq analyses. Bone marrow mononuclear cells (BMNCs) isolated as above from HSC-xenografted mice were labeled with our reference antibody panel. hu-CD45^+^ cell were first partitioned based on CD19 and CD10 before subdivision of double negative CD34hi HSPCs into CD45RA^-/int^ or CD45RA^hi^ cellular fractions; CD45RA^hi^ cells were further subdivided into LMDPs and CD127^-^ or CD127^+^ compartments using adjoining gates.

**Extended Data Fig. 2.**
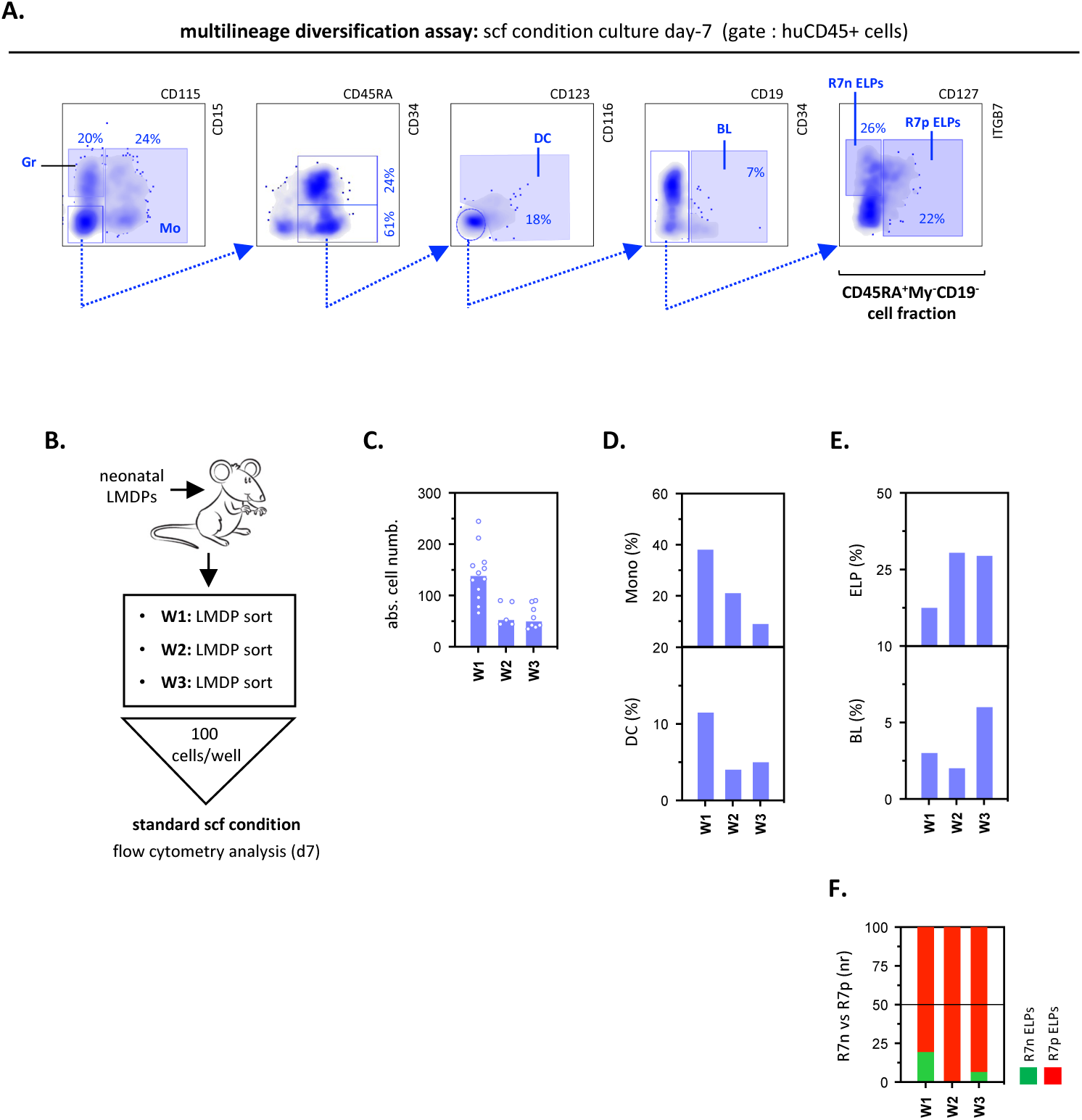
Assessment of lineage output in bulk diversification assays. (A). Gating strategy used for the quantification of the lympho-myeloid output in multilineage diversification assays. In this setting, LMDPs isolated from the BM HSC-xenografted mice were seeded by 100 cell-pools in 96-well culture plates and cultured for 7 days onto OP9 cells under the standard *scf* condition. Analysis by flow cytometry was performed using a 13-antibody panel. After exclusion of doublets, gates were set on hu-CD45^+^ cells which were first divided into CD115^-^CD15^+^ granulocytic (Gr) or CD115^+^CD15^-^ monocytic (Mo) subsets. Remaining CD45RA^+^ (CD115^-^CD15^-^) cells were further partitioned based on CD116 and CD123 to capture CD123^+^CD116^-/+^ dendritic cells (DCs). The CD123^-^CD116^-^ cellular fraction was fractionated into CD19^+^ (BLs) or CD19^-^ fractions. Further subdivision of the CD19^-^ compartment according to differential ITGB7 and CD127 expression allowed isolation of the CD127^-^ (R7^-^) or CD127^+^ (R7^+^) ELPs. Blue arrows indicate the gating procedure; bi-dimensional density plots showing expression of the indicated markers are from ≥ 10 concatenated wells. Percentages of the corresponding cellular subsets are indicated. Results are representative of > 50 experiments. Of note, the same gating procedure was applied for the characterization of single-cell derived clones. (B-F). Time-dependent variation in lympho-myeloid potentials of LMDPs isolated from LMDP-xenografted mice. (B) Experimental design: mice xenografted with neonatal LMDPs were sacrificed at the indicated endpoints before assessment of LMDP lympho-myeloid potential in bulk diversification assays. LMDPs were cultured by 100 cell-pools for 7 days under the standard *scf* condition before quantification by flow cytometry of absolute cell yields and lineage outputs. Bar plots show (C) absolute cell yields (bars indicate median cell numbers; circles correspond to individual wells); median percentages of (D) CD115^+^ monocytes (top) and CD123^+^ DC (bottom) or of lymphoid (E) ELPs (top) and BLs (bottom); results are normalized relative to total hu-CD45^+^ cells); (F) Stacked bar plots showing normalized ratios of CD127^-^ (green) *vs*. CD127^+^ (red) ELPs. Results expressed as median percentages of 5 to ≥ 20 replicates are from one representative experiment out of 2.

**Extended data Fig. 3.**
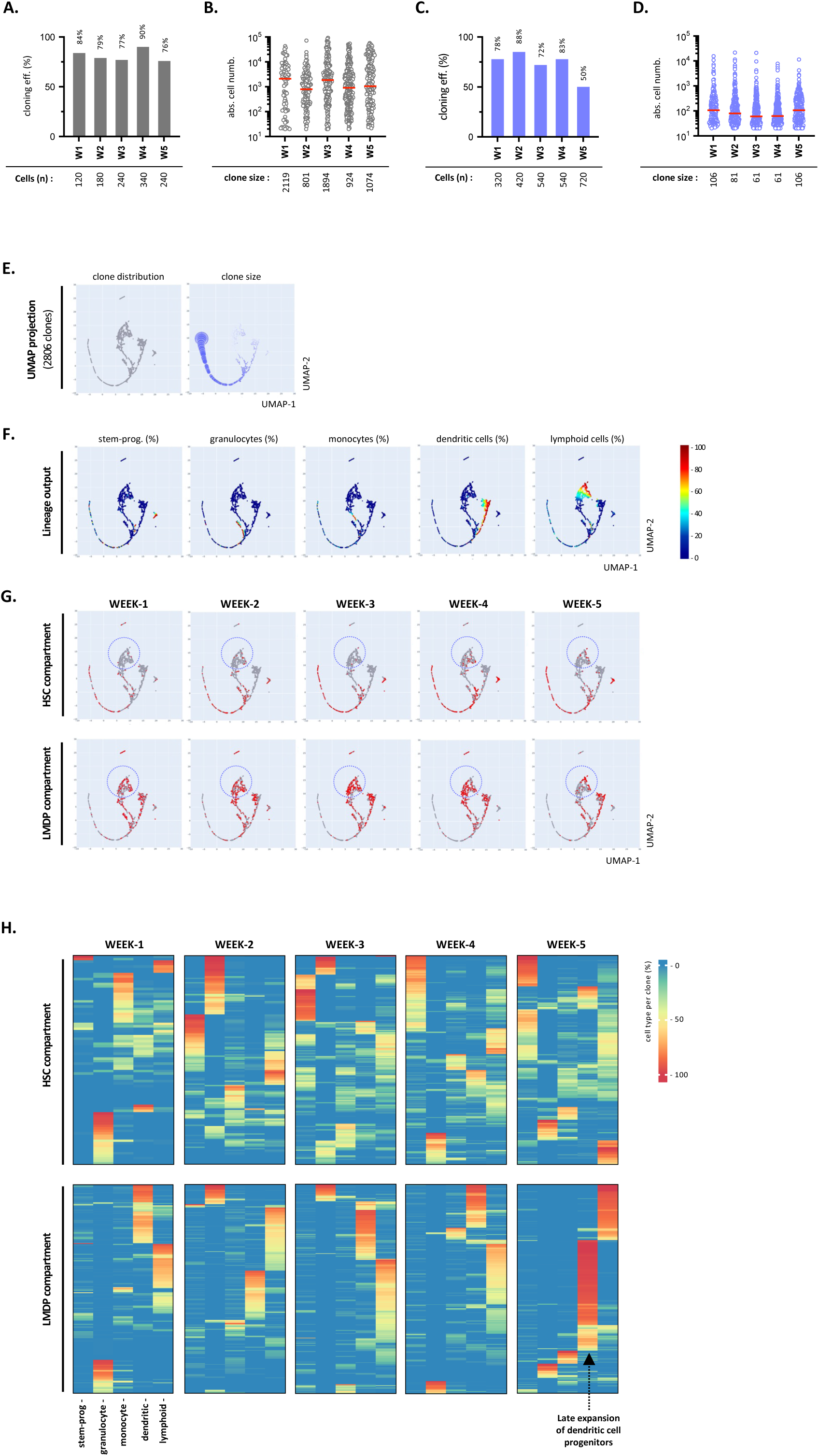
Assessment of lineage output in clonal diversification assays. Mice xenografted with neonatal HSCs were sacrificed at weekly endpoints before assessment of the lympho-myeloid potential of single HSCs or LMDPs in clonal diversification assays. Cells were seeded and cultured individually for 14 days under the *scf:gm-csf:tpo* condition before quantification by flow cytometry of absolute cell yields and lineage outputs (see also Extended Data Fig. 1A for the gating procedure); positivity threshold for clone detection was set arbitrarily at ≥ 20 cells/clone. Percentages of the different cell types are normalized relative to total hu-CD45^+^ cells; lymphoid cell percentages are defined as the sum of the percentages of CD127^-^ or CD127^+^ ELPs and CD19^+^ B-cells. (A-D) Shown are cloning efficiencies and median clone sizes of HSC- or LMDP-derived clones. (A, B) Bar plots show the cloning efficiency of the corresponding (A) HSCs (gray) or (C) LMDPs (blue); total numbers of seeded cells are indicated. Circle plots show absolute cell numbers per (B) HSC- or (D) LMDP-derived clone; red bars indicate medians; corresponding values are indicated (lower row). Results are pooled from 3 independent experiments. (E-G). Application of UMAP algorithm to data pooled from all time points and ancestor cells (n=2806). (E) dot (left panel) and bubble (right panel) maps show the projection of all ancestor cells onto the UMAP coordinates; bubbles indicate clone sizes; (F) heatmaps show the projection of stem/progenitor cells, as well as of granulocytic, monocytic, dendritic or lymphoid potentials; (G) snapshots show the projection onto the UMAP of clones derived from HSC (upper panel) or LMDP (lower panel) ancestors sorted at the indicated time points; dashed circles correspond to the projection area of lymphoid-enriched clones. (H) Heatmaps show hierarchical clustering of clones derived from individual HSCs (upper HM) or LMDPs (lower HM) isolated at the indicated time points; clones are stratified according to lineage output; hierarchical clustering is based on euclidian distance.

**Extended Data Fig. 4.**
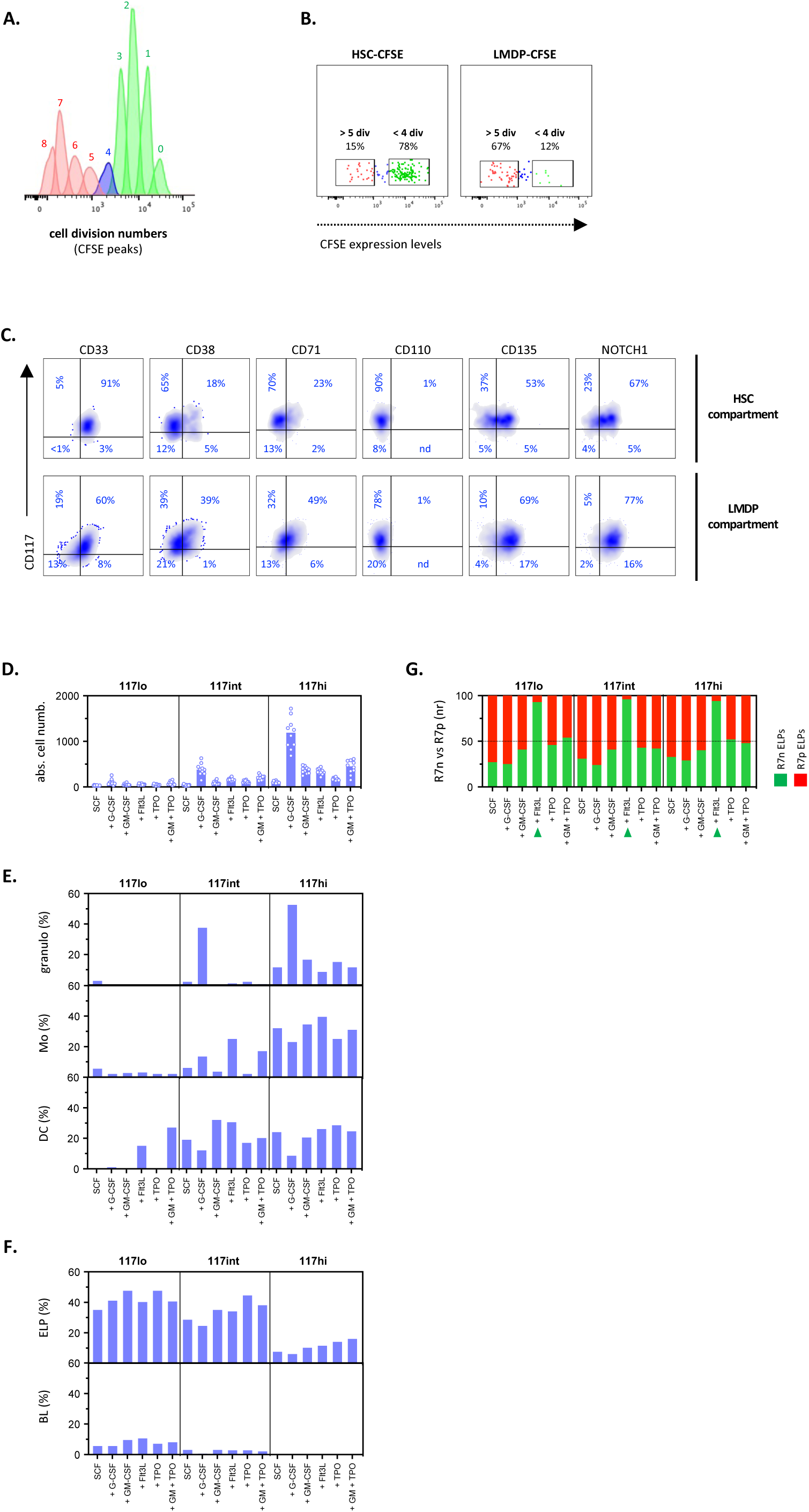
Functional characterization of CD34^hi^ HSPC fractions. (A, B) Fractionation of BM HSCs or LMDPs based on the number of conservative divisions. Neonatal HSCs or LMDPs were labeled with proliferation-dependent dye carboxyfluorescein diacetate succinimidyl ester (CFSE) prior injection to irradiated NSG mice (2.5 x 10^5^ cells/mouse). At day 8 after grafting hu-CD45^+^ BM cells were labeled with a 13-antibody panel before subdivision of self-renewed HSCs or LMDPs according to the number of conservative cell divisions (<4 *versus* >5 divisions). Corresponding divided versus non-divided HSCs or LMDPs were seeded individually in diversification assays and cultured for 14-days under clonal scf:gm-csf:tpo condition. Assessment of clone size and quantification lineage output was performed by flow cytometry. (A). Determination of the cell division numbers of BM HSCs or LMDPs based on “FlowJo proliferation tool” (peak 0 correspond to undivided cells). (B). Gating strategy to subdivide the HSC or LMDP compartments into “divided” (> 5 divisions) or “non-divided” (< 4 divisions) fractions; percentages of cells are indicated. (C). Immunophenotypic substratification of the LMDP compartment. Hu-CD45^+^ cells harvested at week-3 after grafting from the BM of mice reconstituted with neonatal HSCs were labeled with 13 to 15 antibody panels. Density plots show expression of corresponding markers within the HSC (upper panel) or LMDP (lower panel) compartments; percentages are labeled cells indicated. (D-G). Effect of growth/differentiation factors on the lymphoid-myeloid output of CD117^high-int-low^ LMDP fractions. CD117^high, int, low^ LMDPs subdivided as shown in Figure 3A were seeded for 7 days by 100 cell-pools in diversification assays under the standard SCF condition supplemented or not with the indicated growth factors before quantification of absolute cell yields and lympho-myeloid utputs. (D) Shown are absolute cell yields; bars indicate median cell numbers; circles correspond to individual wells. (E-F) Bar plots showing median percentages of (E) myeloid granulocytes (Granulo; upper panel), monocytes (Mo; medium panel) and dendritic cells (DC; lower panel) or of (F) lymphoid CD127^-/+^ ELPs (upper panel) and BLs (lower panel) outputs. (G) Stacked bar plots show normalized ratios of CD127^-^ (green) vs. CD127^+^ (red) ELPs; results are expressed as median percentages of 10 replicates pooled from a representative experiment.

**Extended Data Fig. 5.**
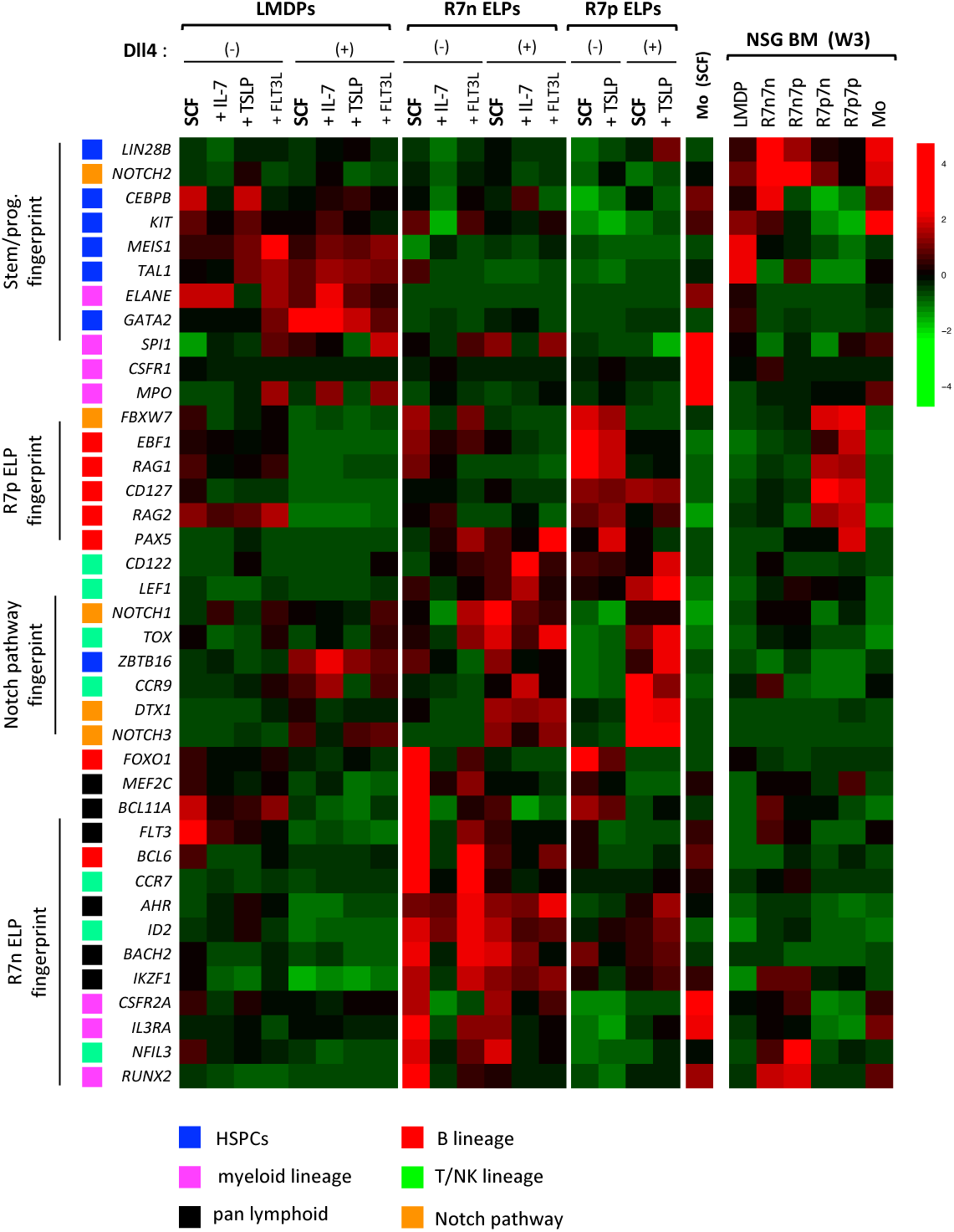
Lineage fingerprints of in vitro-generated LMDPs, CD127^-^ (R7n) or CD127^+^ (R7p) ELPs and CD115^+^ Mo. LMDPs isolated from the BM of HSC-engrafted were seeded as above and cultured as above onto OP9 or OP9-DL4 cells with *scf* condition in the presence or absence of IL-7, TSLP and Flt3L. At culture day-7, the indicated cell subsets (LMDPs, CD127^-^ [R7n] or CD127^+^ [R7n] ELPs, CD115^+^ Mo) were sorted and processed by multiplex PCR for expression (40 genes; primer list is provided as Table S2A). Primary LMDPs, CD127^-^CD7^-/+^ (R7n7n or R7n7p) or CD127^+^CD7^-/+^ (R7p7n or R7p7p) ELP and CD115^+^ Mo sorted from the BM of W3 NSG-UCB mice were used as reference for lineage fingerprints (*see also Alhaj Hussen et al. Immunity 2017; 47:680; DOI: 10.1016/j.immuni.2017.09.009*). Hierarchical clustering is based on 39 genes expressed by 25 cell subsets. Gene expression levels are normalized relative to HPRT. Data are means of 4-6 biological replicates. Colored squares show lineage-specific fingerprints

**Extended Data Fig. 6.**
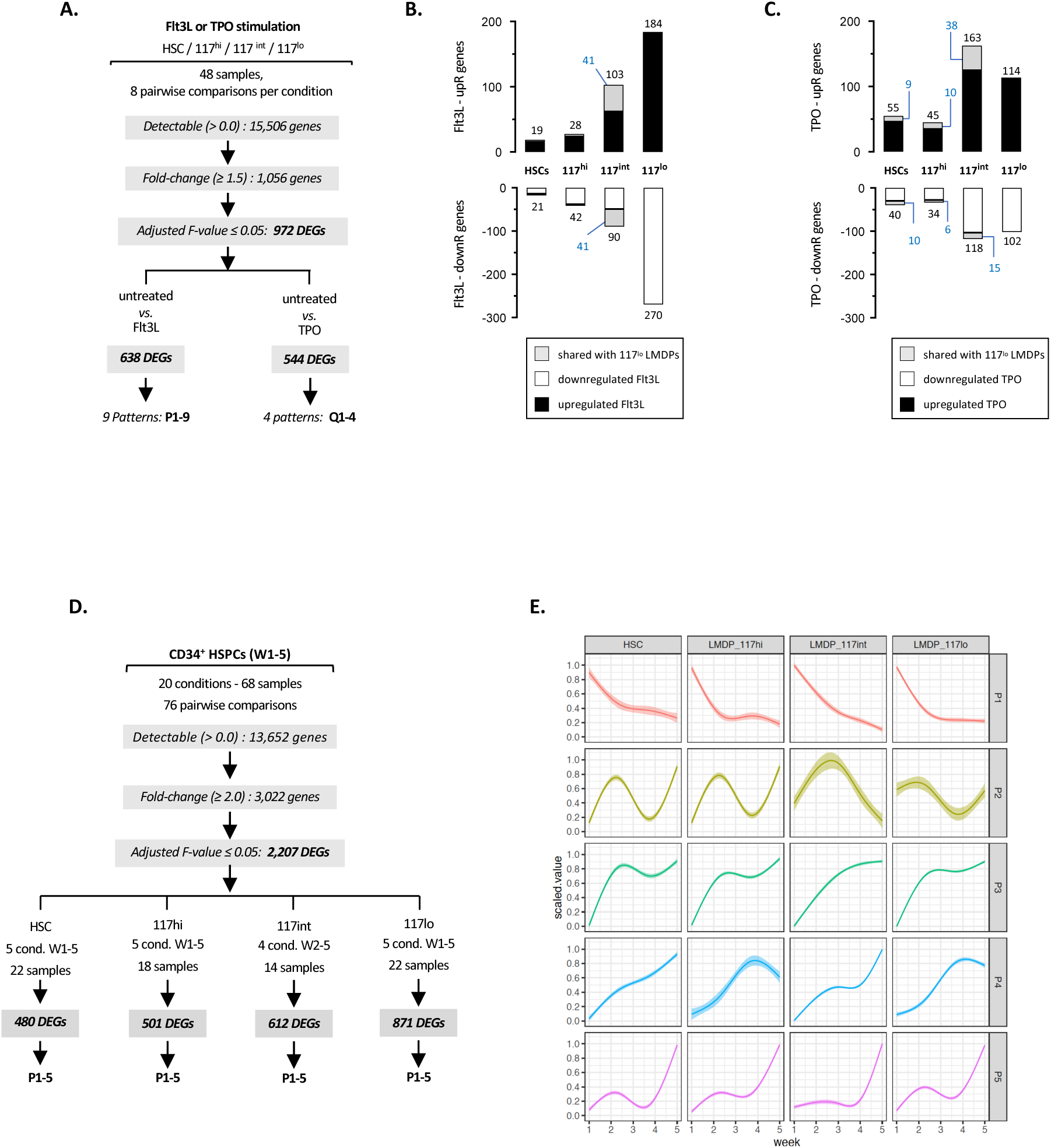
Mini-RNA sequencing analyses. (A-C). Transcriptional response of CD34^+^ HSPCs to short-term stimulation with Flt3L or TPO. The indicated HSC, CD117^hi-int^ LMDP and CD117^lo^ MLP subsets were seeded by 500 cell pools in 96-well plates and cultured for 6 hours under the scf condition with or without Flt3L or TPO (10 ng/ml each) before processing for mini-RNAseq. (A) Flowchart summarizing the filtration and clustering strategies used for extraction of population-specific gene signatures across the corresponding cellular subsets and culture conditions; detection thresholds, fold-changes and adjusted F-value are indicated. Barplots showing the absolute numbers of (B) Flt3L- or (C) TPO-responsive genes. Black and empty bars correspond respectively of up- or downmodulated genes; gray bars and blue numbers correspond to genes shared between CD117^lo^ MLPs and the other cell subsets. (D, E). Time-dependent variation in gene expression patterns across 4 HSC, CD117^hi-int^ LMDP and CD117^lo^ MLP subsets. HSC-xenografted mice were sacrificed at the indicated time points before sorting of the indicated subsets and processing for mini-RNAseq. (D) Pipeline based on pairwise comparisons used for extraction of population- and time-specific gene signatures across the indicated time points and cellular subsets; detection thresholds, fold-changes and adjusted F-value are indicated. (E) Dynamic trends of 2207 DEGs distributed across 5 temporal patterns shared between the indicated subsets. Results are log-transformed, normalized and scaled. Since at week-1 after grafting LMDPs express homogeneously high CD117 they were not further fractionated (unf: unfractionated); also, due to limited cell yields, week-2 LMDPs were subdivided into CD117^hi^ LMDP or CD117^lo^ MLP fractions.

**Extended Data Fig. 7.**
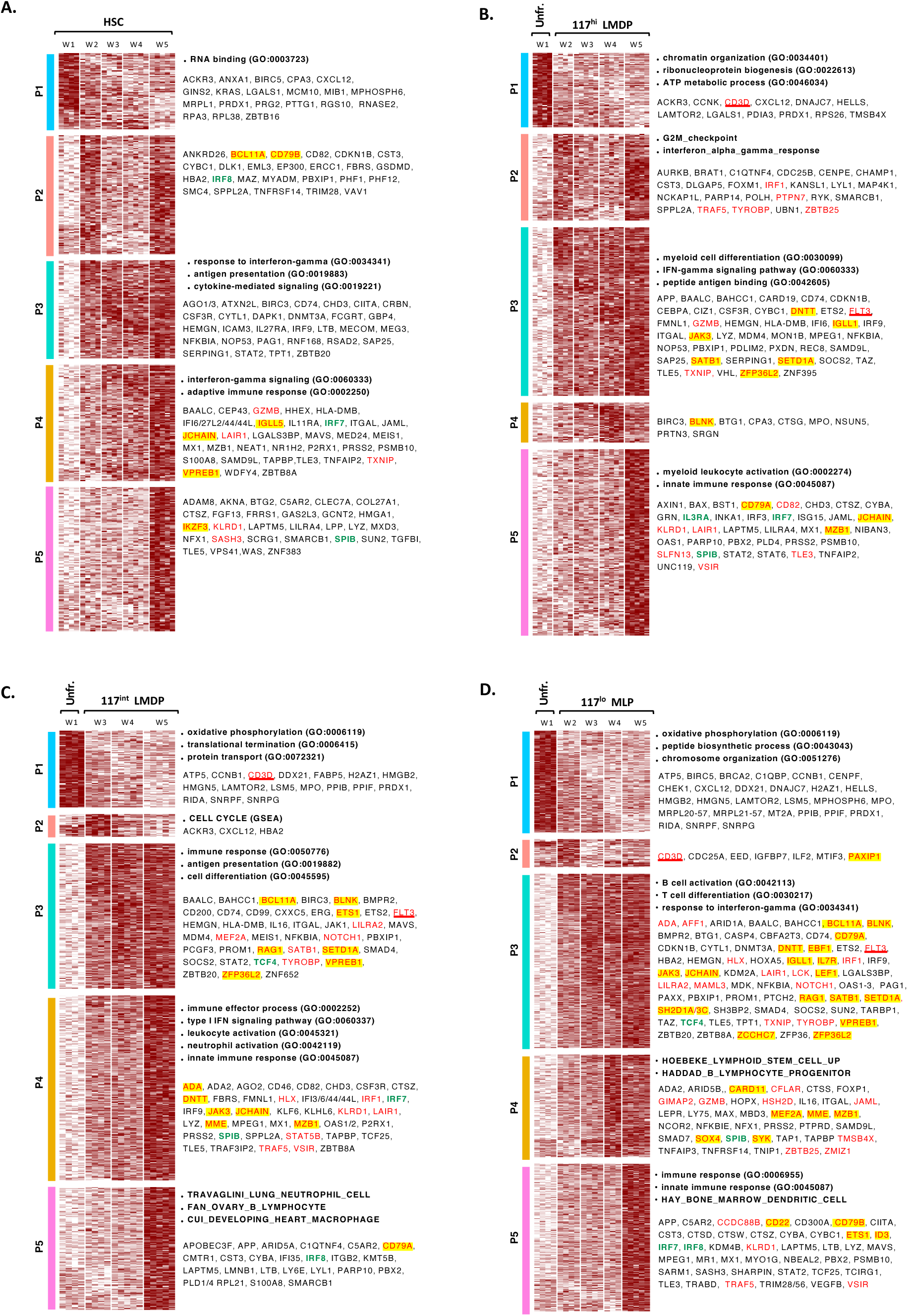
Time-dependent variation in HSC and CD117^hi, int, lo^ LMDP gene signatures. Heatmaps showing time-dependent variation in expression of 2207 DEGs distributed across 5 clusters across the indicated (A) HSC or (B) CD117^hi^ LMDP, (C) CD117^int^ LMDP or (D) CD117^lo^ MLP compartments. Gene expression values are log-transformed, normalized and scaled. Genes linked to lymphoid (red) or dendritic (bold green) differentiation are indicated; B-lymphoid genes are highlighted in yellow; CD3D and FLT3 genes are underlined (bold red).

**Extended Data Fig. 8.**
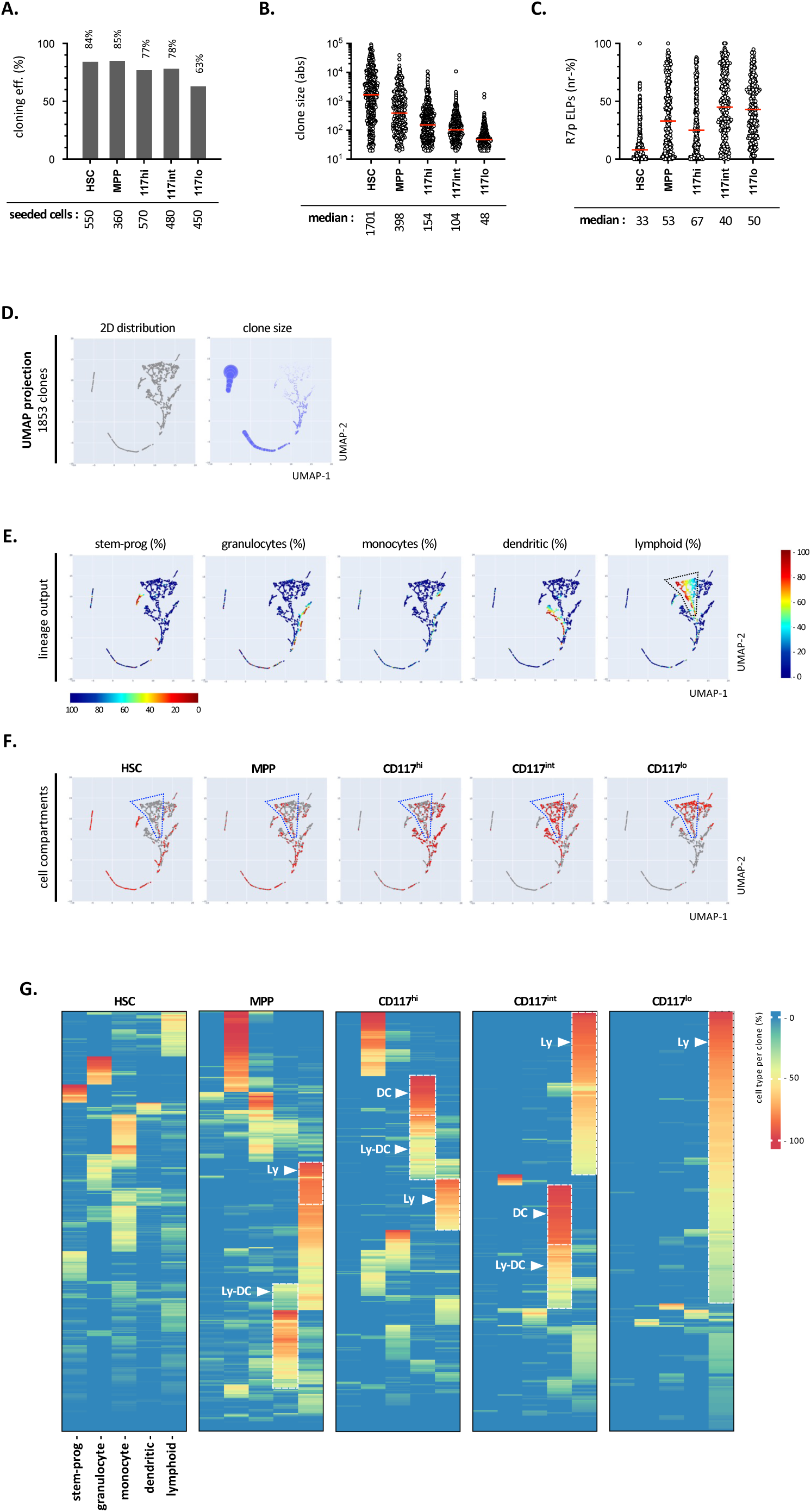
Simultaneous assessment of the clonal architecture of the indicated cellular ccompartments. The indicated HSC, MPP, CD117^hi-int^ LMDP and CD117^lo^ MLP subsets (HSC-xenografted mice; week-3) were individually seeded and cultured for 14 days under the clonal *scf:gm-csf:tpo* condition before quantification of lympho-myeloid output (see also legend of Figure 2). Results are pooled from a representative experiment. (A) Shown are the cloning efficiencies of the indicated subsets; total numbers of seeded cells are indicated. (B, C) Circle plots showing the (B) absolute cell numbers per clone or (C) relative percentages of CD127^+^ ELPs normalized relative to total ELP content on a per clone basis (positivity threshold for ELP detection is set arbitrarily at ≥ 10 ELPs/clone); red bars indicate medians; corresponding values are indicated (lower row). Assessment of statistical significance was performed with the Mann-Whitney test (**** p<0.0001; *** p<0.001; ** p<0.01;). Results are pooled from 1 representative experiment. (D-F). UMAP projection of data pooled from all HSC, MPP, CD117^hi-int^ LMDP and CD117^lo^ MLP ancestor cells (n=1853) : (D) dot (left) and bubble show projection of the compendium of ancestor cells onto the UMAP coordinates; diameters of bubbles indicate clone size; (E) heatmap showing the projection of relative percentages of CD34^+^ stem/progenitor, CD15^+^ granulocytic, CD115^+^ monocytic, CD123^+^ dendritic or of lymphoid cells on a per clone basis; (F) Snapshots show the projection of the indicated HSC, MPP or CD117^hig-int-low^ LMDP ancestor cells; dashed polygon indicate the projection area of lymphoid clones. (G). Heatmaps showing the lineage output of clones derived from individual HSCs, MPPs, CD117^hi-int^ LMDPs and CD117^lo^ MLPs. Results are expressed as normalized percentages of the indicated cell subsets; hierarchical clustering uses Euclidian distance. Lymphoid cell percentages are defined as the sum of the percentages of CD19^+^ BLs and CD127^-/+^ ELPs. Dashed lines show clones with selective enrichment in lymphoid, dendritic or mixed lympho-dendritic potential.

**Extended Data Fig. 9.**
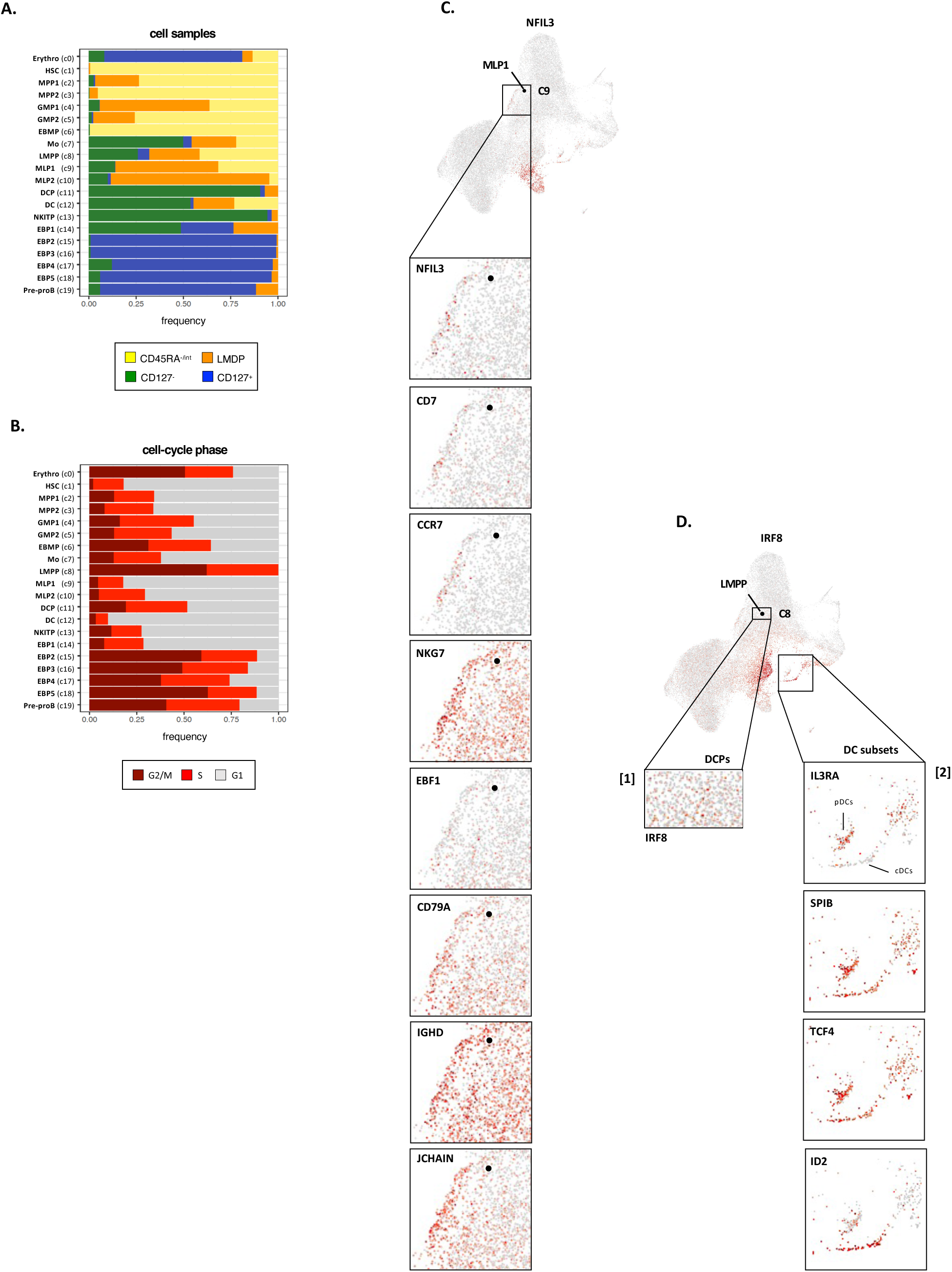
Single-cell RNA-sequencing analyses. (A, B). Repartition of clusters C0-19 according to immunophenotype or cell-cycle status. The indicated CD45RA^-/int^; CD45RA^hi^ and CD127^-^ or CD127^+^ HSPC compartments were isolated from the BM of HSC-xenografted mice (week-3) and subjected to scRNA-seq profiling with the 10X genomic platform. (A) Stacked bar plot showing the relative distribution of cluster-specific cells between LMDP, CD127^-^ or CD127^+^ compartments. As expected, MLP clusters (C9-10) mainly distribute within the LMDPs, whereas NKITPs (C12) and DCPs (C13) are exlusively detected amongst CD127^-^ HPCs. Most immature EBP1s that start upregulating IL7R/CD127 are situated at the interface between LMDPs, CD127^-^ and CD127^+^ HPCs, while downstream EBP2-5 and proB cells fall exlusively within the CD127^+^ compartment. (B) Stacked bar plot showing the distribution of cluster-specific cells according to cell-cycle status; relative proportions of cluster-specific cells are indicated. Note that whereas LMPPs are in G2/M or S phases, a majority of MLPs (C9-10) stand in G1 phase, consistent with the idea that lymphoid commitment is associated with a proliferation arrest. Further, in constrast to EBP1s that do not proliferate, EBP2-5 and preproB cells are actively cycling. (C, D). UMAP visualization of lineage-specific genes. (C) Single-cell mapping of the direct MLP differentiation pathway: inserts show expression of the indicated lymphoid marker genes in individual cell projecting in the immediate vicinity of MLP2 (C9). (D) Single-cell mapping of DC differentiation pathways; inserts show expression of [1] dendritic TF *IRF8* within the C8 LMPP cluster or [2] of established markers of plasmacytoid (pDC) (*IL3R*, *SPIB*, *TCF4*) or conventional (cDC) (*ID2*) dendritic cells.

**Extended Data Fig. 10.**
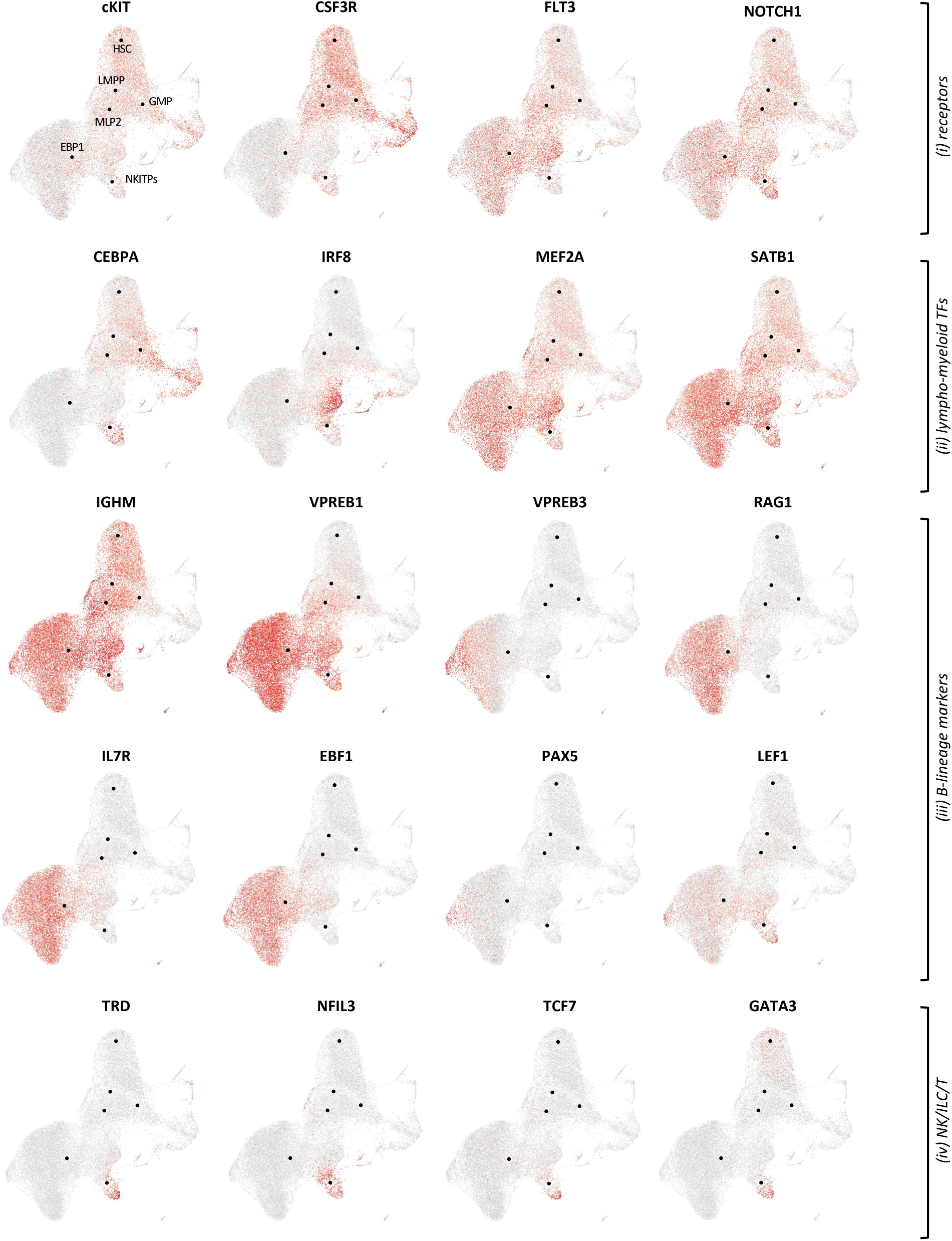
Single-cell RNA-sequencing analyses (*continued*). Shown is expression in individual cells of 20 genes corresponding to (*i*) receptors of growth/differentiation factors (*cKIT*, *Flt3*, *CSF3R*, *Notch1*), (*ii*) TFs regulating lympho-myeloid specifcation (*CEBPA*, *IRF8, SATB1*, *MEF2A*, *LEF1*), (iii) B-lymphoid genes (*IGHM*, *VPREB1*, *VPREB3*, *RAG1, IL7R, EBF1*, *PAX5*); (i*v*) NK/ILC/T lineages genes (*TRD*, *NFIL3*, *TCF7*, *GATA3*). Black dots indicate the medoids of corresponding HSC (C1), LMPP (C8), MLP2 (C10), GMP1 (C4), EBP1 (C14) and NKITP (C13) clusters.

**Extended Data Fig. 11.**
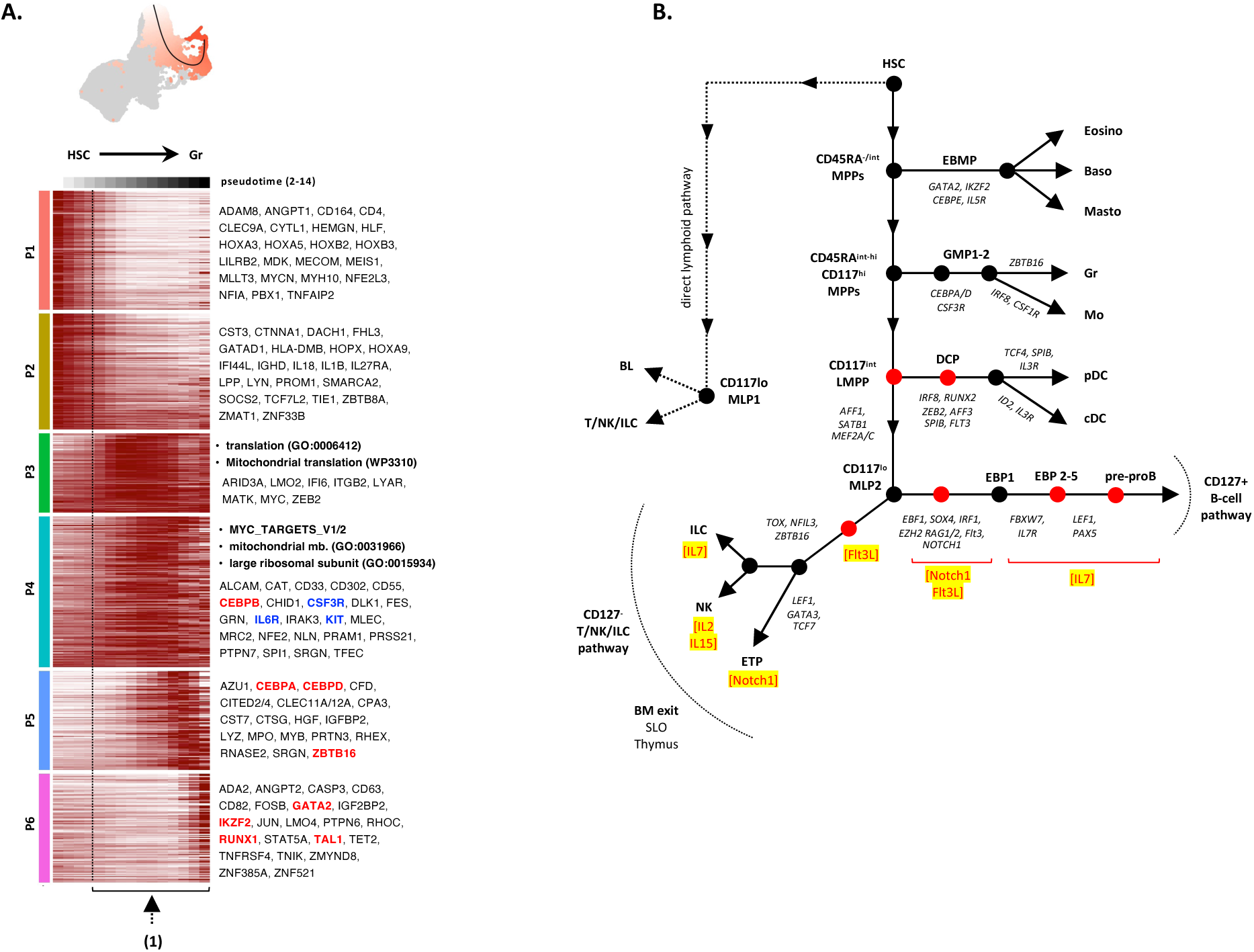
Single-cell RNA-sequencing analyses *(continued-2)* and proposed model of hematpoietic organization. (A). dendritic lineage differentiation trajectory. UMAP shows the granulocytic differentiation trajectory represented as smoothed black curve; lineage-specific cells are color-coded (white to red) according to pseudotime values. Lower heatmap represents min to max normalized expression of dynamically variable genes; trajectory inference analyses were performed with SlingShot. Growth factor receptors are in blue; TFs are in red. Dashed line and dashed arrow indicate cell proliferation. (B). An integrated model of lymphoid architecture. Lymphoid specification follows distinct direct (MLP1) or stepwise (MLP2) differentiation pathways converging towards the generation of CD117^lo^ MLPs. Differentiation along the direct pathway (dashed lines; left side) proceeds in absence of multi-lineage diversification, progression along the stepwise MLP2 pathway (plain lines; right side) follows a classical hierarchy where multi-lineage diversification starts downstream of HSCs with sequential emergence of EBMPs and then GMPs. Following individualization of GMP1s, CD45RA^hi^ MPPs start downmodulating CD117 to differentiate into CD117^int^ LMPPs. At this stage cells undergo a proliferative burst during which they acquire cell-fate biases resulting in the subsequent generation of bipotent lympho-dendritic or unipotent lymphoid or dendritic progenitors. Downstream lymphoid commitment of CD117^lo^ MLPs is associated with a cell proliferation arrest that also marks the first lymphoid development checkpoint. Subsequent choice between B (CD127^+^) and T/NK/ILC (CD127^-^) lymphoid pathways relies on a dynamic equilibrium between the intrinsic B-lineage bias of CD117^lo^ MLPs and extrinsic Flt3-signaling that drives their differentiation along the T/NK/ILC lymphoid pathway. Downstream progression along the CD127^+^ B-lymphoid pathway comprises a first phase of amplification controlled by Notch1 and Flt3L driving differentiation into EBP1s whose proliferation arrest coincides with downmodulation of *NOTCH1* and *FLT3* as and upregulation of transcripts coding Notch-inhibitor *FBXW7* and *IL-7R*. Downstream EBP2-5 subsets undergo a second phase of IL7-dependent amplification driving their progression toward pre-proB cells that again with stop proliferating and initiate IgH locus recombination as they acquire surface CD10/CD24/CD19. Differentiation along the T/NK/ILC pathway comprises a single wave of Flt3L-dependent proliferation allowing individualization of lineage-specified NK, ILC or T precursors that will exit the BM to reach the thymus or secondary lymphoid organs to complete their differentiation into mature effectors. NK precursors become strictly dependent on IL-2/15 for subsequent maturation and proliferation, while ILC progenitors upregulate IL-7R before diversifying into ILC1, ILC2 or ILC3 precursors. Lastly, ETPs leave the BM to reach the thymus. Following establishment of intra-thymic lympho-stromal synapses, Notch1- and to a lesser extent Flt3-signaling synergistically promote ETP proliferation, IL7R upregulation and drive T-lineage commitment followed by onset of TCRδ/β recombination. It has to be noted that this revised model differs from the previously proposed “2-family model” of lymmhoid architecture (*Alhaj Hussen et al. Immunity 2017; 47:680; DOI: 10.1016/j.immuni.2017.09.009*) by taking into account the dynamics of *IL7R* expression along the T/NK/ILC or B lymphoid pathways. EBMP: eosinophil/basophil/mast cell progenitor; GMP: granulo-monocyte progenitor; MPP: multipotent progeniors; LMPP: lymphoid-primed multipotent progenitor; DCP: dendritic cell progenitor; MLP: multi-lymphoid progenitor; ELP: early lymphoid progenitor; EBP: early B cell precursor; NK: natural killer; ILC: innate lymphoid cells; ETP: early T-cell progenitor. Black circles indicate key differentiation stages and development checkpoints; red circles correspond to cell proliferation phases. Key signaling pathways (red) are highlighted (yellow).

## REFERENCES

1. Laurenti, E. & Gottgens, B. From haematopoietic stem cells to complex differentiation landscapes. Nature 553, 418–426 (2018).

2. Pronk, C.J. et al. Elucidation of the phenotypic, functional, and molecular topography of a myeloerythroid progenitor cell hierarchy. Cell stem cell 1, 428–442 (2007).

3. Sanjuan-Pla, A. et al. Platelet-biased stem cells reside at the apex of the haematopoietic stem-cell hierarchy. Nature 502, 232–236 (2013).

4. Yamamoto, R. et al. Clonal analysis unveils self-renewing lineage-restricted progenitors generated directly from hematopoietic stem cells. Cell 154, 1112–1126 (2013).

5. Mossadegh-Keller, N. et al. M-CSF instructs myeloid lineage fate in single haematopoietic stem cells. Nature 497, 239–243 (2013).

6. Rieger, M.A., Hoppe, P.S., Smejkal, B.M., Eitelhuber, A.C. & Schroeder, T. Hematopoietic cytokines can instruct lineage choice. Science 325, 217–218 (2009).

7. Benz, C. et al. Hematopoietic stem cell subtypes expand differentially during development and display distinct lymphopoietic programs. Cell stem cell 10, 273–283 (2012).

8. Rodriguez-Fraticelli, A.E. et al. Clonal analysis of lineage fate in native haematopoiesis. Nature 553, 212–216 (2018).

9. Kim, S. et al. Dynamics of HSPC repopulation in nonhuman primates revealed by a decade-long clonal-tracking study. Cell stem cell 14, 473–485 (2014).

10. Biasco, L. et al. In Vivo Tracking of Human Hematopoiesis Reveals Patterns of Clonal Dynamics during Early and Steady-State Reconstitution Phases. Cell stem cell 19, 107–119 (2016).

11. Six, E. et al. Clonal tracking in gene therapy patients reveals a diversity of human hematopoietic differentiation programs. Blood 135, 1219–1231 (2020).

12. Weinreb, C., Rodriguez-Fraticelli, A., Camargo, F.D. & Klein, A.M. Lineage tracing on transcriptional landscapes links state to fate during differentiation. Science 367 (2020).

13. Park, B.G. et al. Reconstitution of lymphocyte subpopulations after hematopoietic stem cell transplantation: comparison of hematologic malignancies and donor types in event-free patients. Leuk Res 39, 1334–1341 (2015).

14. Upadhaya, S. et al. Kinetics of adult hematopoietic stem cell differentiation in vivo. J Exp Med 215, 2815–2832 (2018).

15. Adolfsson, J. et al. Identification of Flt3+ lympho-myeloid stem cells lacking erythro-megakaryocytic potential a revised road map for adult blood lineage commitment. Cell 121, 295–306 (2005).

16. Sitnicka, E. et al. Key role of flt3 ligand in regulation of the common lymphoid progenitor but not in maintenance of the hematopoietic stem cell pool. Immunity 17, 463–472 (2002).

17. Amann-Zalcenstein, D. et al. A new lymphoid-primed progenitor marked by Dach1 downregulation identified with single cell multi-omics. Nat Immunol 21, 1574–1584 (2020).

18. Igarashi, H., Gregory, S.C., Yokota, T., Sakaguchi, N. & Kincade, P.W. Transcription from the RAG1 locus marks the earliest lymphocyte progenitors in bone marrow. Immunity 17, 117–130 (2002).

19. Klein, F. et al. Dntt expression reveals developmental hierarchy and lineage specification of hematopoietic progenitors. Nat Immunol 23, 505–517 (2022).

20. Cumano, A. et al. New Molecular Insights into Immune Cell Development. Annu Rev Immunol 37, 497–519 (2019).

21. Canque, B. et al. Characterization of dendritic cell differentiation pathways from cord blood CD34(+)CD7(+)CD45RA(+) hematopoietic progenitor cells. Blood 96, 3748–3756 (2000).

22. Fritsch, G. et al. Rapid discrimination of early CD34+ myeloid progenitors using CD45-RA analysis. Blood 81, 2301–2309 (1993).

23. Doulatov, S. et al. Revised map of the human progenitor hierarchy shows the origin of macrophages and dendritic cells in early lymphoid development. Nat Immunol 11, 585–593 (2010).

24. Haddad, R. et al. Molecular characterization of early human T/NK and B-lymphoid progenitor cells in umbilical cord blood. Blood 104, 3918–3926 (2004).

25. Hao, Q.L. et al. Identification of a novel, human multilymphoid progenitor in cord blood. Blood 97, 3683–3690 (2001).

26. Hoebeke, I. et al. T-, B- and NK-lymphoid, but not myeloid cells arise from human CD34(+)CD38(-)CD7(+) common lymphoid progenitors expressing lymphoid-specific genes. Leukemia 21, 311–319 (2007).

27. Galy, A., Travis, M., Cen, D. & Chen, B. Human T, B, natural killer, and dendritic cells arise from a common bone marrow progenitor cell subset. Immunity 3, 459–473 (1995).

28. Karamitros, D. et al. Single-cell analysis reveals the continuum of human lympho-myeloid progenitor cells. Nat Immunol 19, 85–97 (2018).

29. Haddad, R. et al. Dynamics of thymus-colonizing cells during human development. Immunity 24, 217–230 (2006).

30. Parietti, V. et al. Dynamics of human prothymocytes and xenogeneic thymopoiesis in hematopoietic stem cell-engrafted nonobese diabetic-SCID/IL-2rgammanull mice. J Immunol 189, 1648–1660 (2012).

31. Alhaj Hussen, K. et al. Molecular and Functional Characterization of Lymphoid Progenitor Subsets Reveals a Bipartite Architecture of Human Lymphopoiesis. Immunity 47, 680–696 e688 (2017).

32. Lin, C.C. et al. Knock-out of Hopx disrupts stemness and quiescence of hematopoietic stem cells in mice. Oncogene 39, 5112–5123 (2020).

33. Vitali, C. et al. SOCS2 Controls Proliferation and Stemness of Hematopoietic Cells under Stress Conditions and Its Deregulation Marks Unfavorable Acute Leukemias. Cancer Res 75, 2387–2399 (2015).

34. Mann, Z., Sengar, M., Verma, Y.K., Rajalingam, R. & Raghav, P.K. Hematopoietic Stem Cell Factors: Their Functional Role in Self-Renewal and Clinical Aspects. Front Cell Dev Biol 10, 664261 (2022).

35. de Laval, B. et al. Thrombopoietin-increased DNA-PK-dependent DNA repair limits hematopoietic stem and progenitor cell mutagenesis in response to DNA damage. Cell stem cell 12, 37–48 (2013).

36. Roy, A. et al. Transitions in lineage specification and gene regulatory networks in hematopoietic stem/progenitor cells over human development. Cell Rep 36, 109698 (2021).

37. Street, K. et al. Slingshot: cell lineage and pseudotime inference for single-cell transcriptomics. BMC genomics 19, 477 (2018).

38. Velten, L. et al. Human haematopoietic stem cell lineage commitment is a continuous process. Nat Cell Biol 19, 271–281 (2017).

39. Mansson, R. et al. Molecular evidence for hierarchical transcriptional lineage priming in fetal and adult stem cells and multipotent progenitors. Immunity 26, 407–419 (2007).

40. Alexander, T.B. et al. The genetic basis and cell of origin of mixed phenotype acute leukaemia. Nature 562, 373–379 (2018).

41. Allman, D. et al. Thymopoiesis independent of common lymphoid progenitors. Nat Immunol 4, 168–174 (2003).

42. Berthault, C. et al. Asynchronous lineage priming determines commitment to T cell and B cell lineages in fetal liver. Nat Immunol 18, 1139–1149 (2017).

43. Kawamoto, H., Ikawa, T., Ohmura, K., Fujimoto, S. & Katsura, Y. T cell progenitors emerge earlier than B cell progenitors in the murine fetal liver. Immunity 12, 441–450 (2000).

44. Sitnicka, E. et al. Complementary signaling through flt3 and interleukin-7 receptor alpha is indispensable for fetal and adult B cell genesis. J Exp Med 198, 1495–1506 (2003).

45. Sitnicka, E. et al. Critical role of FLT3 ligand in IL-7 receptor independent T lymphopoiesis and regulation of lymphoid-primed multipotent progenitors. Blood 110, 2955–2964 (2007).

46. Paul, F. et al. Transcriptional Heterogeneity and Lineage Commitment in Myeloid Progenitors. Cell 164, 325 (2016).

47. Cacchiarelli, D. et al. Integrative Analyses of Human Reprogramming Reveal Dynamic Nature of Induced Pluripotency. Cell 162, 412–424 (2015).

48. Thielecke, L. et al. Limitations and challenges of genetic barcode quantification. Sci Rep 7, 43249 (2017).

49. Thielecke, L., Cornils, K. & Glauche, I. genBaRcode: a comprehensive R-package for genetic barcode analysis. Bioinformatics 36, 2189–2194 (2020).

50. Becht, E. et al. Dimensionality reduction for visualizing single-cell data using UMAP. Nat Biotechnol (2018).

51. Giacosa, S. et al. Cooperative Blockade of CK2 and ATM Kinases Drives Apoptosis in VHL-Deficient Renal Carcinoma Cells through ROS Overproduction. Cancers (Basel) 13 (2021).

52. Love, M.I., Huber, W. & Anders, S. Moderated estimation of fold change and dispersion for RNA-seq data with DESeq2. Genome biology 15, 550 (2014).

53. Chalmel, F. & Primig, M. The Annotation, Mapping, Expression and Network (AMEN) suite of tools for molecular systems biology. BMC Bioinformatics 9, 86 (2008).

54. Ritchie, J.B., Tovar, D.A. & Carlson, T.A. Emerging Object Representations in the Visual System Predict Reaction Times for Categorization. PLoS Comput Biol 11, e1004316 (2015).

55. Smyth, G.K. Linear models and empirical bayes methods for assessing differential expression in microarray experiments. Stat Appl Genet Mol Biol 3, Article3 (2004).

56. Ritchie, M.E. et al. limma powers differential expression analyses for RNA-sequencing and microarray studies. Nucleic acids research 43, e47 (2015).

57. Wettenhall, J.M. & Smyth, G.K. limmaGUI: a graphical user interface for linear modeling of microarray data. Bioinformatics 20, 3705–3706 (2004).

58. Van den Berge, K. et al. Trajectory-based differential expression analysis for single-cell sequencing data. Nat Commun 11, 1201 (2020).

